# Increased hippocampal excitability and altered learning dynamics mediate cognitive mapping deficits in human aging

**DOI:** 10.1101/581108

**Authors:** Nadine Diersch, Jose P Valdes-Herrera, Claus Tempelmann, Thomas Wolbers

**Affiliations:** Aging & Cognition Research Group, German Center for Neurodegenerative Diseases (DZNE), Magdeburg, 39120, Germany; Department of Neurology, Otto-von-Guericke University Magdeburg, 39120, Germany; Center for Behavioural Brain Sciences (CBBS), Otto-von-Guericke University Magdeburg, 39120, Germany

**Keywords:** aging, Bayesian modeling, effective connectivity, fMRI, learning, novelty, older adults, spatial navigation, virtual reality

## Abstract

Learning the spatial layout of a novel environment is associated with dynamic activity changes in the hippocampus and in medial parietal areas. With advancing age, the ability to learn spatial environments deteriorates substantially but the underlying neural mechanisms are unknown. Here, we report findings from a behavioral and a fMRI experiment where older and younger adults performed a spatial learning task in a photorealistic virtual environment. We modeled individual learning states using a Bayesian state-space model and found that activity in retrosplenial cortex/parieto-occipital sulcus and anterior hippocampus did not change systematically as a function learning in older compared to younger adults across repeated episodes in the environment. Moreover, effective connectivity analyses revealed that the age-related learning deficits are linked to an increase in hippocampal excitability. Together, these results provide important insights into how human aging affects computations in the brain’s navigation system, highlighting the critical role of the hippocampus.

## INTRODUCTION

Venturing into the unknown and exploring our surroundings has always been one of the hallmarks of human identity. To do so, we must be able to quickly encode novel environments, generate a cognitive map of them, and flexibly retrieve this information later. This allows us to navigate effortlessly in an ever-changing world. With advancing age, however, these abilities deteriorate considerably (Lester, Moffat, Wiener, Barnes, & Wolbers, 2017). Older adults have been found to be slower in learning novel environments and have problems to retrieve this newly learnt information later, even when the results are controlled for their slower learning rate (Iaria, Palermo, Committeri, & Barton, 2009). Moreover, they have problems in learning landmark locations during exploratory navigation (Yamamoto & DeGirolamo, 2012), whereas their route and landmark location memory is relatively preserved for familiar environments (Merriman, Ondřej, Roudaia, O’Sullivan, & Newell, 2016; Rosenbaum, Winocur, Binns, & Moscovitch, 2012). As a consequence, older adults may avoid unfamiliar places and become easily overwhelmed when confronted with changes in their environment.

Although core regions of the brain’s navigation circuit in the medial temporal lobe are among the first to be affected during the progression from healthy aging to Alzheimer’s disease (AD; Braak & Del Tredici, 2015), the neural mechanisms for age-related deficits in spatial learning are still poorly understood, even in healthy older adults. Studies in rodents and non-human primates showed that place cells in the CA3 subfield of the hippocampus exhibit higher firing rates in aged animals during navigation, and they fail to encode new information when rats encounter novel environments (Thomé, Gray, Erickson, Lipa, & Barnes, 2016; Wilson, Ikonen, Gallagher, Eichenbaum, & Tanila, 2005). Moreover, firing patterns of aged CA1 place cells are often unstable across repeated visits to the same environment (Barnes, Suster, Shen, & McNaughton, 1997; see also Hok, Chah, Reilly, & O’Mara, 2012). In humans, in contrast, there is evidence for an age-related activity reduction in the hippocampus and medial parietal areas during spatial navigation (Konishi et al., 2013; Moffat, Elkins, & Resnick, 2006). This might be related to an age-related shift from hippocampal-dependent allocentric navigational strategies to egocentric strategies that are associated with activity in extrahippocampal structures (e.g., caudate), which might be more or less beneficial depending on the specific task demands (cf., Schuck, Doeller, Polk, Lindenberger, & Li, 2015).

However, whether higher or lower activity in certain brain regions is indicative of a compensatory mechanism or neural dedifferentiation in older adults is a long-standing issue in cognitive neuroscience research on aging (Grady, 2012; Park & Reuter-Lorenz, 2009). Evidence from studies investigating age-related impairments in separating sensory input from existing mnemonic representations (i.e., pattern separation) suggests that hyperactivity in hippocampal subfields (dentate gyrus and CA3) may underlie memory deficits in healthy aging (Reagh et al., 2018; Yassa et al., 2011). Hippocampal hyperactivity has been further associated with increased beta-amyloid (Aβ) accumulation, one of the earliest indicators for a progression to AD (Leal, Landau, Bell, & Jagust, 2017), and APOEε4 status, a genetic risk factor for AD (Sinha et al., 2018). At present, however, it is unknown whether this effect also pertains to age-related problems in retrieving the layout of spatial environments.

Age-related differences in neural activity may further depend on the point in time when activity is measured during task performance (e.g., during or after learning). Studies in younger adults showed that the engagement of the retrosplenial cortex (RSC) and the parieto-occipital sulcus (POS) together with the hippocampus changes over the course of learning (Auger, Zeidman, & Maguire, 2015; Brodt et al., 2016; Patai et al., 2019; Wolbers & Büchel, 2005). In Wolbers and Büchel (2005), for example, participants were repeatedly transported through a novel virtual environment containing multiple landmarks, and they were subsequently asked to retrieve the spatial position of the landmarks in relation to each other. While activity in the RSC/POS tracked learning of the relative landmark locations and increased with increasing performance across learning sessions, hippocampal activity reflected the amount of learning in a given session and decreased after participants performed at ceiling. Moreover, the functional coupling between RSC/POS and hippocampus is increased as long as the hippocampus is involved during the process of learning (Auger et al., 2015; Brodt et al., 2016).

Given the time course of its involvement during spatial learning, the RSC has been implicated in the storage and retrieval of hippocampal-dependent spatial and episodic memories. It has reciprocal interactions with a multitude of cortical and subcortical brain regions (Bzdok et al., 2015; Kobayashi & Amaral, 2003, 2007). It receives inputs from CA1 and the subiculum and is known to be involved in the integration of different spatial reference frames/viewpoints as well as in updating spatial representations in response to novel sensory input (Epstein, 2008; Miller, Vedder, Law, & Smith, 2014; Mitchell, Czajkowski, Zhang, Jeffery, & Nelson, 2018). The hippocampus, in turn, particularly its anterior portion, is known for its role in generating (spatial) representations in response to sensory input and prior experience (Zeidman & Maguire, 2016). For example, place-cell like activity in the RSC of mice, which develops gradually during spatial learning, critically relies on intact input from the hippocampus to support memory retrieval, and it does not evolve in animals with hippocampal lesions (Mao et al., 2018). Thus, age-related problems in retrieving newly learnt information during spatial navigation might point to a malfunctioning of the integration of hippocampal input within RSC and/or a corrupted hippocampal signal.

Here, we report findings from two experiments where we 1) characterized age-related problems in learning a novel virtual environment (VE), by measuring how well healthy older and younger adults are able to retrieve its spatial layout across several learning blocks, and 2) investigated the underlying neural mechanisms using fMRI. We focused on activity changes in the RSC/POS and the hippocampus and changes in effective connectivity within and between the two regions. We modeled spatial learning using a Bayesian implementation of a state-space model, because behavioral performance is noisy and may not accurately reflect the subject-specific learning state. We used the outputs of the model in the analysis of the fMRI data to examine intra-and inter-individual differences in learning. Behaviorally, we found that although some older adults learned just as well as younger adults, the majority of them showed substantial deficits in retrieving information about the initially unfamiliar environment. Importantly, while we replicated findings from earlier studies in younger adults, we found that activity in the RSC/POS and the anterior hippocampus did not change systematically in older adults across repeated episodes in the environment. We additionally found that activity in the posterior hippocampus reflected inter-individual differences in learning within older adults. Finally, we show that their learning deficits are linked to impaired spatial information processing in the anterior hippocampus, as evidenced by an age-related increase in the intrinsic excitability of this region during task performance.

## RESULTS

Findings are reported from two separate samples comprising healthy younger and older adults who performed a spatial learning task either purely behaviorally (17 younger adults and 17 older adults) or in a combined fMRI-behavioral experiment (25 younger adults and 32 older adults). In both experiments, following an initial familiarization phase before testing/outside of the scanner, eight learning blocks were implemented during which eight retrieval phases alternated with seven encoding phases (see Figure 1A for the detailed structure of the fMRI experiment). The experiments were implemented using a complex VE resembling a realistic city center of a typical German town, which was unfamiliar to our participants (Figure 1B). During retrieval, participants’ knowledge about the spatial layout of the VE was tested. They were asked to point in direction of one of two target landmarks while being located in the middle of one of four intersections that were approached from one of four directions, while the background was obscured by fog (Figure 1C, Video 2). In the behavioral experiment, participants performed 96 navigational retrieval trials (12 per learning block) in a pseudo-randomized order, with the restriction that each intersection/target landmark combination was encountered from two of the four possible directions in the first half of the experiment. In the second half of the experiment, the intersections were approached from the remaining two directions. This allowed us to examine how experiencing familiar locations from a novel viewpoint affects pointing performance. In the fMRI experiment, 64 navigational retrieval trials were implemented in a randomized order (8 per learning block). The retrieval phases additionally contained 32 control trials (4 per learning block) consisting of a color discrimination task that did not require the retrieval of the spatial layout of the VE. During encoding phases, which did not differ between experiments, participants were passively transported around the whole environment while being instructed to memorize its layout and the positions of the target landmarks in relation to the intersections (see Video 1 for a short exemplary segment of one encoding tour).

**Figure 1.**
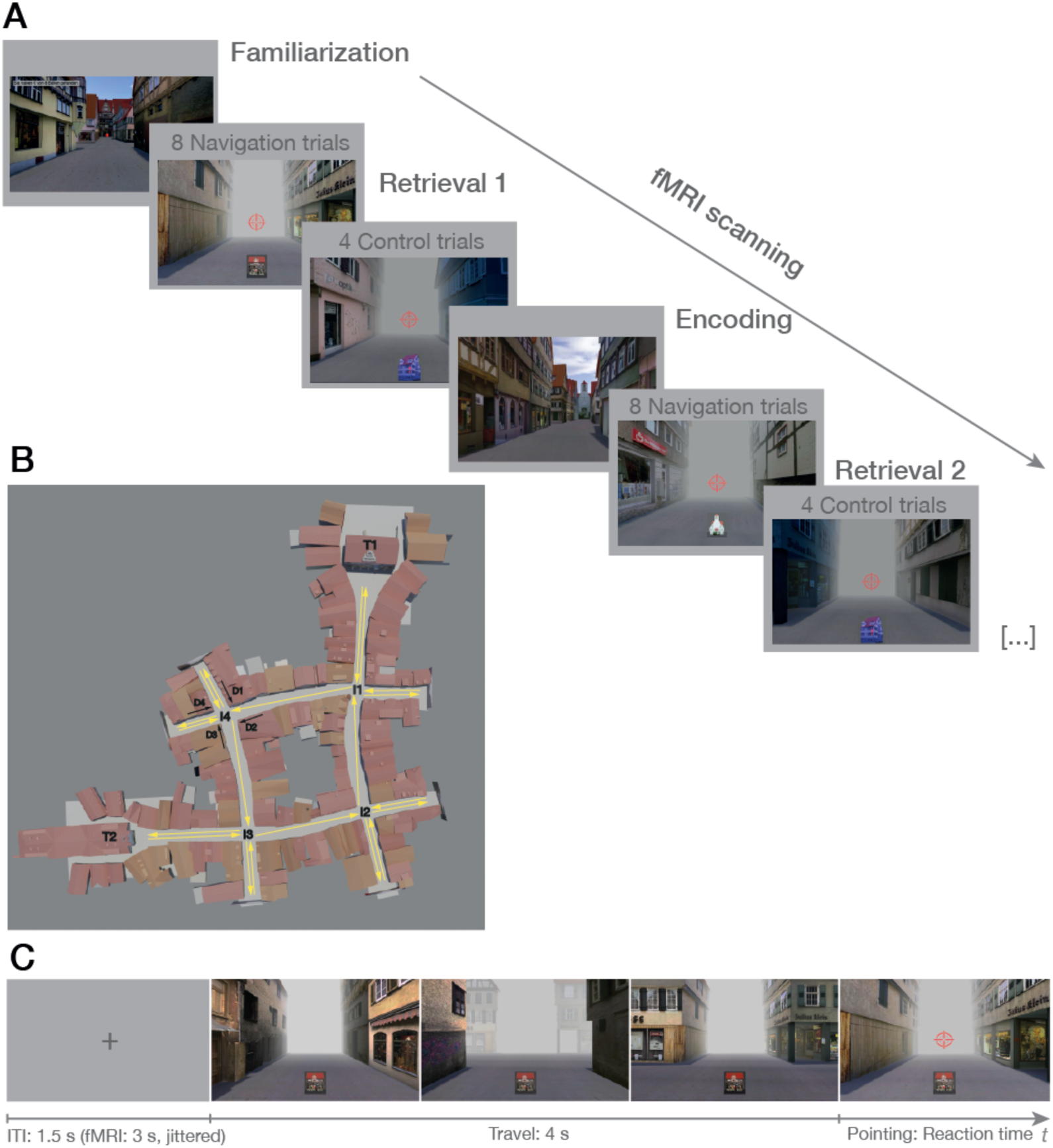
Spatial learning task. (A) Procedure of the fMRI experiment. After a familiarization phase outside of the scanner, eight retrieval phases, each comprising 8 navigational retrieval trials and 4 control trials, alternated with 7 encoding phases during scanning. In the behavioral experiment, the structure was the same except that 12 navigational retrieval trials per learning block were completed while the control trials were omitted. (B) Layout of the virtual environment (VE). The VE resembled a typical German historic city center and consisted of four interconnected intersections (I1-I4) that could be reached from 4 directions (D1-D4). At two intersections, a town hall (T1) and a church (T2) were placed at the end of one of the outgoing streets that served as target landmarks in the navigational retrieval trials. Yellow arrows exemplify one encoding tour that started from one of the target landmarks in clockwise or counterclockwise direction in counterbalanced order across the experiment (a short segment of one encoding tour is shown in Video 1). (C) Structure of one example navigational retrieval trial to measure spatial learning (see also Video 2). After fixation, participants were passively transported towards one of four intersections in the VE starting from one of the four streets leading towards that intersection. Movement stopped at the center of the intersection, a red crosshair appeared, and participants were asked to move the crosshair in direction to the respective target landmark. During the entire duration of the trial, a picture cue of the target landmark was displayed at the bottom of the screen, and the background was obscured by fog. In the fMRI experiment, an additional jittered interval of 1 s (still phase) was added after the travel phase/before the crosshair appeared on screen.

We used the angular deviation of the participants’ response from the respective target landmark (i.e., absolute pointing errors) to measure performance improvements across learning blocks. However, performance in these kinds of tasks can be corrupted by various noise sources and, hence, might not accurately reflect the actual learning state of the participant. Therefore, subject-specific improvements in navigational performance were estimated by using a Bayesian implementation of a state-space model that disambiguated learning from random trial-by-trial fluctuations in performance (see Figure S1 and Bayesian Modeling of Performance Data section; Commandeur & Koopman, 2007). We modeled learning block-wise because learning in our task primarily took place during the encoding phases and not during individual retrieval trials.

### Behavioral experiment

#### Lower performance and reduced learning in older adults

An ANOVA with learning block (1-8) as repeated measures variable and age-group (younger adults, older adults) as between-subjects variable on the average absolute pointing errors showed significant main effects of learning block, F(7, 224) = 19.5, p < .001, and age group, F(1, 32) = 85.2, p < .001. This was modulated by a significant interaction between the two factors, F(7, 224) = 7.40, p < .001. At the beginning, both age groups performed around chance level (90°), but older adults showed lower performance and less improvement over the course of the experiment compared to younger adults (Figure 2A). The change in direction from the first to the second half of the experiment had no major effect on this pattern of results as shown by a non-significant interaction between learning block and age group when directly comparing the fourth and fifth learning block, F(1, 32) = 1.96, p = .171. A separate ANOVA within the older age group on pointing performance per learning block confirmed that older adults generally improved on the task over time as evidenced by a significant main effect of learning, F(7, 112) = 2.58, p = .017. According to the outputs of the Bayesian state-space model (Figure 2C, see also Figure S4A-B for average pointing errors per learning block for each participant), most of the younger adults learned the spatial layout of the VE very fast, reaching ceiling performance already after the first few learning blocks. The older adults, in contrast, differed more widely in their ability to learn.

**Figure 2.**
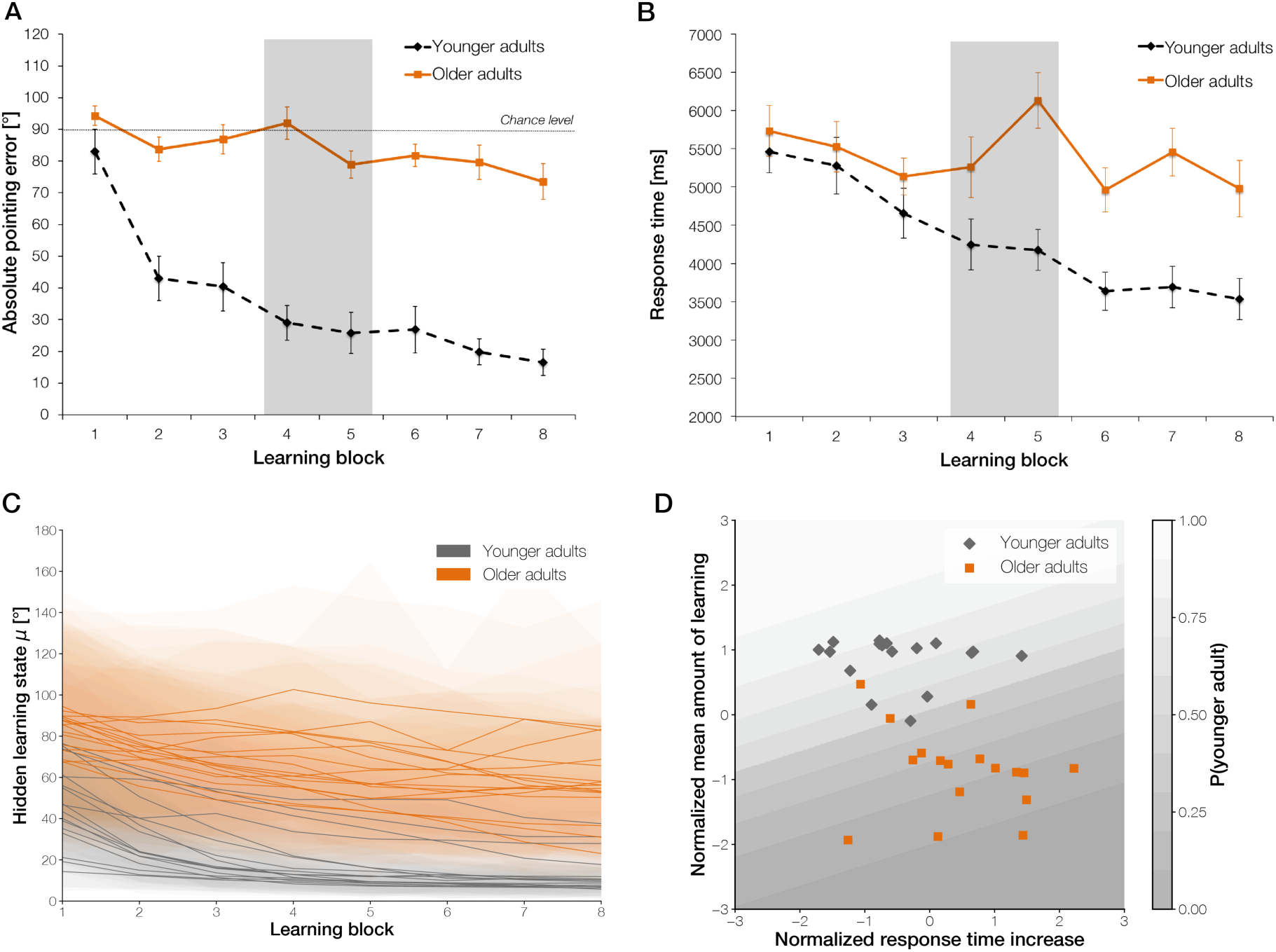
(A) Average absolute pointing errors and (B) response times across the eight learning blocks in older (solid line) and younger adults (dashed line) in the behavioral experiment (highlighted in grey is the 4th and 5th learning block where the change in directions took place from which the intersections were approached). Error bars denote standard errors of the means (SE). (C) Mean estimated performance improvement (hidden learning state) of each participant in the older (orange) and younger (grey) age group, including the standard deviation (SD) of the posterior distributions (shaded area) across learning blocks. (D) Logistic regression results classifying age group membership (shaded lines depict the probability of being classified as a younger adult) based on two behavioral performance features, i.e., the mean amount of learning across the experiment and the increase in response times after the first half of the experiment. See also Figure S4A-B for average pointing errors per learning block for each participant in each age group and Data S1 for additional performance analyses depending on the characteristics of the VE. The Bayesian state-space model to estimate the subject-specific hidden learning states can be found in Figure S1.

To investigate potential biases in pointing behavior that might differ between the age groups, such as an increased tendency to point along streets, circular statistics were applied on the signed pointing error data relative to each target landmark for every intersection-direction combination. From the 32 age-group comparisons (4 intersections × 4 directions × 2 target landmarks), only 7 reached significance as determined by a Watson-Williams test, all p ≤ .047. Older adults showed larger deviation from the correct angle than younger adults in 6 of the 7 instances. The direction of the deviations in pointing (e.g., to the left or right relative to the target landmark), however, varied and none of the effects survived when correcting for multiple comparisons. Thus, we did not find any reliable indications for age-related biases in pointing behavior (see also Data S1 for further analyses of potential performance differences between age groups depending on the characteristics of the VE).

#### Stronger reliance on specific viewpoints in older adults

An ANOVA with learning block (1-8) as repeated measures variable and age group (younger adults, older adults) as between-subjects variable on the response time data confirmed significant main effects of learning block, F(7, 224) = 9.26, p < .001, and age group, F(1, 32) = 10.5, p = .003. Compared to older adults, however, younger adults responded quicker and showed a steeper decline in response times over the course of learning as revealed by a significant interaction between learning block and age group, F(7, 224) = 4.29, p = .001. Notably, when comparing the fourth and fifth learning blocks, a significant interaction between learning block and age group was found, F(1, 32) = 9.34, p = .004. Older but not younger adults showed a substantial increase in response times in the fifth learning block when the intersections were encountered from novel directions (Figure 2B). This result cannot be explained by a confound between pointing performance and required turning at the intersections because the required amount of turning to perform accurately on the task varied from trial to trial depending on the specific intersection-direction-target landmark combination. Moreover, it was kept constant across experiment halves and participants (M = 135°). When considering the fourth and fifth block only, we also did not find any indications that the correct turning angle differed between blocks, age groups, or varied between age groups as a function of learning block, all F ≤ 3.23, p ≥ .082. Thus, older adults’ representations of the spatial layout of the environment seem to be more rigidly tied to the sensory input encountered at the beginning of learning, leading to problems when viewpoints are changing.

#### Individual learning state and reaction time increase after viewpoint change predict age-group

We next used a logistic regression model to check whether age-group can be determined based on two features that characterized age-related performance differences in our task. The mean amount of learning across the whole experiment (i.e., difference between individual learning state estimates from consecutive learning blocks) and the change in response times from the 4th to the 5th learning block served as input features. The model performed very well to estimate the probability of being classified as a younger adult with an average area under the curve (AUC) of 0.99±0.02%. Thus, those participants with a higher probability of belonging to the younger age group show better performance on the task while a higher probability of being in the older age group relates to poorer navigational performance, i.e., a lower mean amount of learning across blocks and a higher increase in response times when previously learned locations are encountered from novel viewpoints (Figure 2D).

### fMRI experiment

After pre-processing of the fMRI data using fmriprep (Esteban et al., 2019) and SPM12, we performed a univariate regression analysis to identify age-related differences in neural activity in the RSC/POS and the hippocampus during different phases of the experiment. We further examined the effects of learning at the within-and between-subject level. Finally, we examined age-and learning-related differences in effective connectivity within and between the two regions.

#### Learning ability varies within the older age group

As in the behavioral experiment, significant main effects for learning block, F(7, 385) = 32.3, p < .001, and age group, F(1, 55) = 167, p < .001, together with a significant interaction between the two factors for the average absolute pointing errors were obtained, F(7, 385) = 11.0, p < .001. This indicates that younger compared to older adults again showed better performance on the task and stronger improvement across learning blocks (Figure 3A). Older adults, however, did show learning at the group level as confirmed by a separate ANOVA within this age group, F(7, 217) = 3.58, p = .001). Accuracy for the control trials was at ceiling across the whole sample (mean proportion of correct responses = 0.97 ± 0.05). The change in directions from the first to the second half of the experiment was omitted here due to the reduced number of trials per learning block as compared to the behavioral experiment. Thus, we did not expect changes in response times from the first to the second half of the experiment.

**Figure 3.**
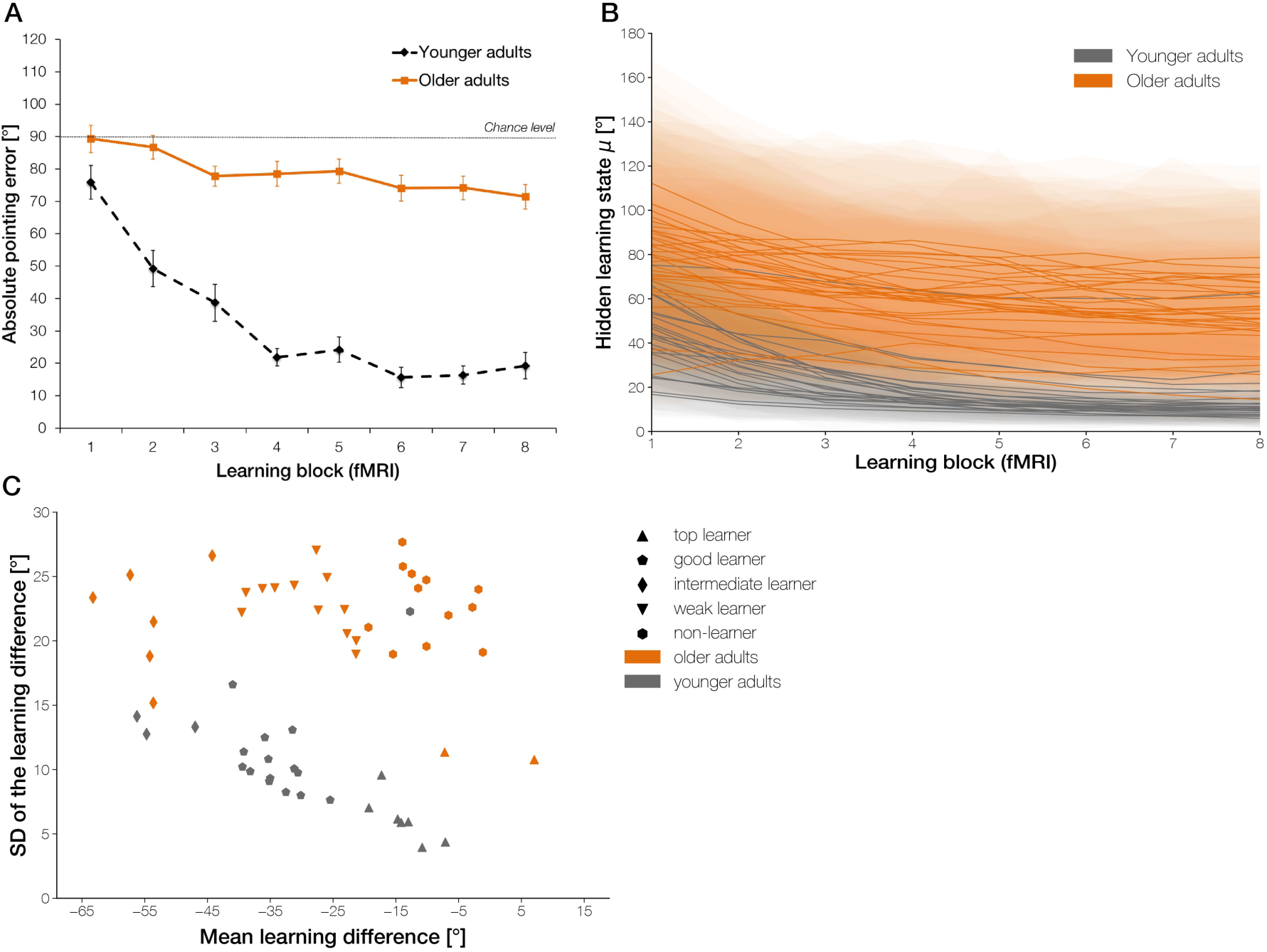
(A) Average absolute pointing errors across the eight learning blocks in older (solid line) and younger adults (dashed line) in the fMRI experiment. Error bars denote standard errors of the means (SE). (B) Mean estimated performance improvement (hidden learning state) of each participant in the older (orange) and younger (grey) age group, including the standard deviation (SD) of the posterior distributions (shaded area) across learning blocks. (C) Learning sub-groups as identified by a K-means clustering algorithm based on the individuals’ overall amount of learning and its SD, as determined by the difference of the latent state distributions of the last and first learning block. See also Figure S4C-D for average pointing errors per learning block for each participant in each age group and S5 for difference distributions, learning state estimates, and performance data for representative individuals from each learning sub-group. The Bayesian state-space model to estimate the subject-specific hidden learning states can be found in Figure S1.

Inspection of the individual learning state estimates as obtained from the state-space model again showed that participants varied substantially in their ability to learn the spatial layout of the VE (Figure 3B, see also Figure S4C-D for average pointing errors per learning block for each participant in each age group). To examine neural activation patterns as a function of the individuals’ overall amount of learning across the experiment, we used a K-means clustering algorithm to identify learning sub-groups based on the difference between the latent state distributions of the last and first learning block (i.e., difference distribution). The estimated optimal number of clusters in our sample turned out to be five (Figure 3C): A group of *top learners* (n = 9), consisting of seven younger adults and two older adults, already learned the layout of the VE after the familiarization phase resulting in a small difference in learning between the first and the last learning block. The second cluster exclusively consisted of younger adults, categorized as *good learners* (n = 14). They typically reached ceiling performance during the first half of the experiment with a low variance in their difference distribution. A group of *intermediate learners* (n = 9), consisting of a three younger and six older adults, were still improving in the second half of the experiment and consequently exhibited the largest difference in their hidden learning state from the beginning to the end of the experiment and a relatively high variance. Individuals belonging to the fourth cluster were categorized as *weak learners* (n = 12) who showed only a small improvement across the whole experiment and a high variance. This cluster consisted exclusively of older adults. Finally, 12 older adults and one younger adult did not show considerable improvement across the learning blocks and were consequently categorized as *non-learners* (n = 13) in the context of our experiment. These between-subject differences in learning demonstrate that our task was not too easy or too difficult for one of the age groups per se. Difference distributions for representative individuals from each learning sub-group, together with learning estimates and behavioral data per learning block, can be found in Figure S5.

#### Activity in hippocampus and RSC/POS is mostly increased in older adults during navigational retrieval

First, to identify age-related differences in activation patterns within the RSC/POS and the hippocampus, irrespective of learning, we contrasted navigational retrieval trials to control trials. Activity in medial parts of the POS/RSC was increased in older compared to younger adults during navigation versus control. This age-related activity increase was also observed in the left anterior hippocampus (Table 1A). An age-related activity reduction was found in a small cluster in the superior right POS and also in a more lateral cluster in the right POS (Table 1B). Second, we checked for the interactions between age group and activation differences during navigational retrieval versus encoding. In one cluster of the right POS as well as two clusters in the right and left anterior hippocampus, activity was increased in older adults compared to younger adults during navigational retrieval versus encoding (Table 1C). There were no clusters within our ROIs where activity was reduced in older adults. Activations outside of our ROIs for these two comparisons and the corresponding results for the whole sample can be found in Table S1A-G.

**Table 1.** Spatial coordinates of the local maxima in the hippocampus and RSC/POS ROI (p < 0.05, FWE-corr.). See Table S1 for the corresponding whole-brain results.

| Brain region | Cluster size | MNI coordinate |  |  | Z-score |
| --- | --- | --- | --- | --- | --- |
|  |  | x | y | z |  |
| A) Increased activity in older compared to younger adults during navigation vs. control |  |  |  |  |  |
| L POS | 83 | -3 | -60 | 34 | 4.56 |
| L RSC |  | -6 | -57 | 21 | 4.32 |
| L Hippocampus | 72 | -21 | -18 | -12 | 4.76 |
|  |  | -30 | -15 | -22 | 4.21 |
| B) Reduced activity in older compared to younger adults during navigation vs. control |  |  |  |  |  |
| R POS | 22 | 12 | -69 | 54 | 4.00 |
|  |  | 18 | -69 | 57 | 3.94 |
|  |  | 15 | -75 | 51 | 3.25 |
| R POS | 55 | 27 | -60 | 24 | 3.96 |
| C) Increased activity in older compared to younger adults during retrieval vs. encoding |  |  |  |  |  |
| R POS | 27 | 12 | -51 | 34 | 4.03 |
| L Hippocampus | 33 | -30 | -15 | -15 | 4.08 |
| R Hippocampus | 20 | 24 | -12 | -12 | 3.97 |
| D) Age-group differences in learning-related activity decreases |  |  |  |  |  |
| R Hippocampus | 20 | 24 | -18 | -15 | 4.55 |
| E) Age-group differences in learning-related activity increases |  |  |  |  |  |
| L POS | 462 | -9 | -66 | 24 | 5.93 |
|  |  | -24 | -72 | 47 | 5.37 |
| L RSC |  | -18 | -57 | 1 | 4.14 |
|  |  | -6 | -63 | 11 | 3.98 |
| R POS | 148 | 24 | -69 | 47 | 4.95 |
|  |  | 21 | -72 | 54 | 4.73 |
| R RSC | 205 | 9 | -57 | 4 | 4.82 |
|  |  | 9 | -63 | 21 | 4.59 |

#### Learning-related activity changes in anterior hippocampus and RSC/POS are less pronounced in older adults

By using the amount of learning per block as contrast weights in our GLM, we assessed age-group differences in the time-course of hippocampal and RSC/POS involvement during navigational retrieval phases. First, we found that activity in the right anterior hippocampus decreased in younger but less so in older adults as a function of learning (Table 1D, Figure 4A). This suggests that hippocampal activity reflected the amount of spatial knowledge that was acquired after each encoding tour in the younger age group. In older adults, in contrast, hippocampal activation did not change systematically across learning blocks. Second, we also found several clusters within the RSC/POS ROI where activity increased over the course of the experiment more in younger than in older adults (Table 1E, Figure 4B). This concerned the whole extent of the left POS from its superior parts to the left RSC, a cluster in the right RSC/POS, and a more lateral cluster in the right POS. Activity in these clusters therefore paralleled performance improvements from learning block to learning block in the younger age group. The older adults’ individual learning curves, in contrast, were again decoupled from activity changes in these regions. Activations outside of our ROIs for these comparisons and the corresponding results for the whole sample can be found in Table S1H-I.

**Figure 4.**
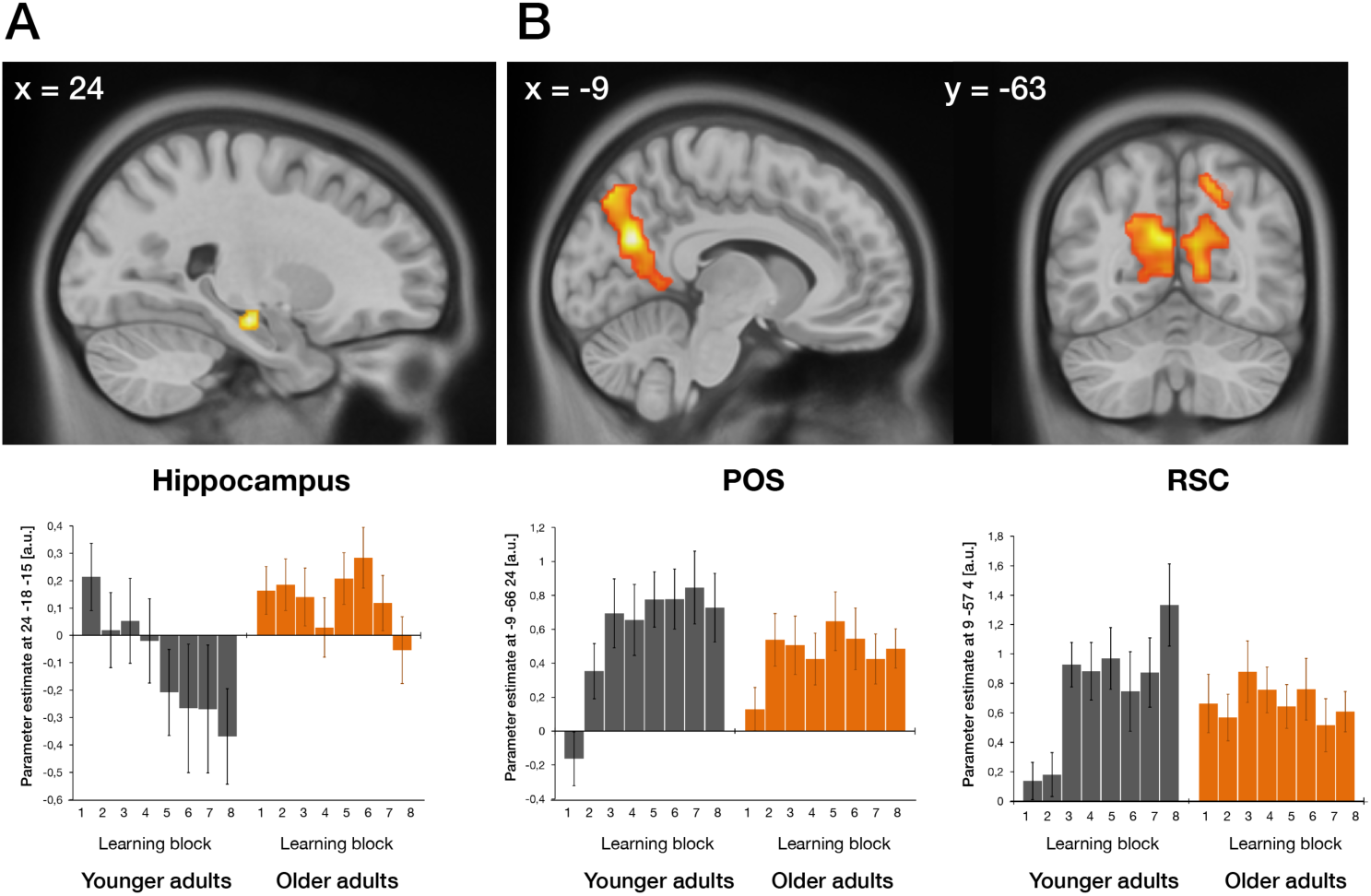
Age-related differences in activity changes in relation to the amount of learning per block during navigational retrieval. Age group differences in (A) hippocampal activity decreases and (B) RSC/POS activity increases across the experiment. Activations are displayed on the 2009 nonlinear asymmetric MNI template that was used for normalization (p < 0.05, FWE-corrected for the respective ROI). Plots depict average parameter estimates of the respective peak voxels per learning block in selected clusters for each age group. Error bars indicate the across-subject standard error of the mean. See also Table S1 for significantly activated clusters elsewhere in the brain.

#### Inter-individual differences in learning within older adults are related to differences in posterior hippocampal engagement

We next included the individual’s learning sub-group as covariate in the analysis to examine in which regions learning-related activity changes across blocks differed as a function of the overall learning ability of the individual. In younger adults, no activations emerged within our ROIs or elsewhere in the brain. However, in older adults, activity in the right posterior hippocampus (24, −30, −9, Z = 5.24, 17 voxels) was differently modulated across learning blocks as a function of the individual’s overall learning ability. Inspection of each sub-group’s parameter estimates revealed that this portion of the hippocampus was generally more involved in top learners as compared to the remaining learning sub-groups (Figure 5). The whole-brain results for this comparison can be found in Table S1J. One should note, however, that there were only two top learners, six intermediate learners and 12 weak as well as 12 non-learners in the older age group and that no activations survived our correction for multiple comparisons within our ROIs or across the whole-brain when testing for the interactions between age group and learning sub-group.

**Figure 5.**
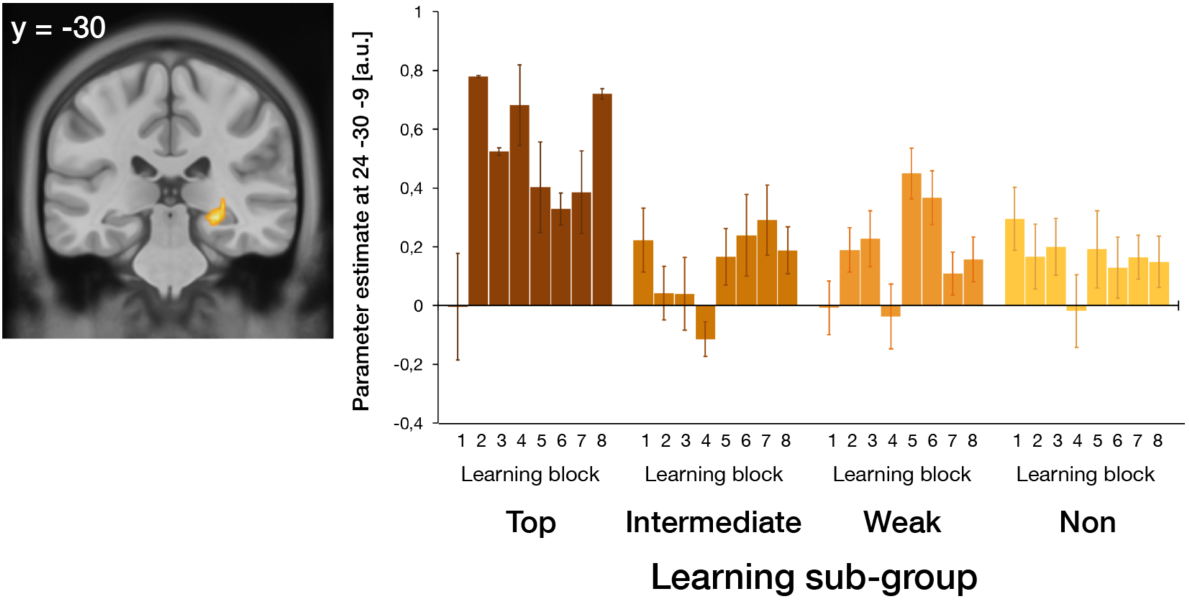
Differential activity changes in relation to the amount of learning per block during navigational retrieval between learning sub-groups in the older age group within the hippocampus. Activations are displayed on the 2009 nonlinear asymmetric MNI template that was used for normalization (p < 0.05, FWE-corrected for the respective ROI). The plot depicts average parameter estimates of the hippocampal peak voxel per learning block for each older learning sub-group. Error bars indicate the across-subject standard error of the mean. See also Table S1 for significantly activated clusters elsewhere in the brain.

#### Age-related reduction in the inhibitory self-connection of the anterior hippocampus

To check whether age-related problems in spatial learning are related to changes in the intrinsic excitability of the anterior hippocampus and the POS/RSC or in the coupling between the two regions, we used the parametric empirical Bayesian (PEB) approach in the context of Dynamic Causal Modeling (DCM; Friston et al., 2016). DCM has been successfully used to determine effective connectivity changes in the hippocampus and related regions during memory processing (Gluth, Sommer, Rieskamp, & Büchel, 2015). Moreover, DCM PEB offers several advantages over classical DCM variants in terms of model selection and second-level group comparisons. First, instead of specifying several models at the first level and comparing their evidence, a full model is estimated for each participant incorporating all parameters of interest, and Bayesian model reduction (BMR) is performed to obtain posterior estimates of nested models in which parameters that do not contribute to the model evidence are pruned. Second, first-level DCMs are equipped with empirical priors that shrink parameter estimates towards a group mean. In this way, each subject’s contribution to the group PEB result is weighted by their precision. Third, applying classical inference methods to examine whether certain parameters differ between groups after model inversion ignores within-subject uncertainty (i.e., variance of the posterior distributions). This is circumvented in PEB by using the full posterior density over the parameters from each participant’s DCM to draw inferences about group level effects.

For each participant, we first specified and estimated a DCM between the right anterior hippocampus and the left POS. Navigational retrieval phases were modeled as driving input into the network via the POS. The amount of learning per block was modeled as modulatory input on the bidirectional connections between the two regions (Figure 6A). In the second-level PEB model, we included age group, learning sub-group, and their interaction as covariates to determine their relative influence on the connection strengths. The left panels in Figure 6B and 6D show the group mean of the average connection strength before and after Bayesian model reduction (BMR), indicating that all four parameters were necessary to explain our data.

**Figure 6.**
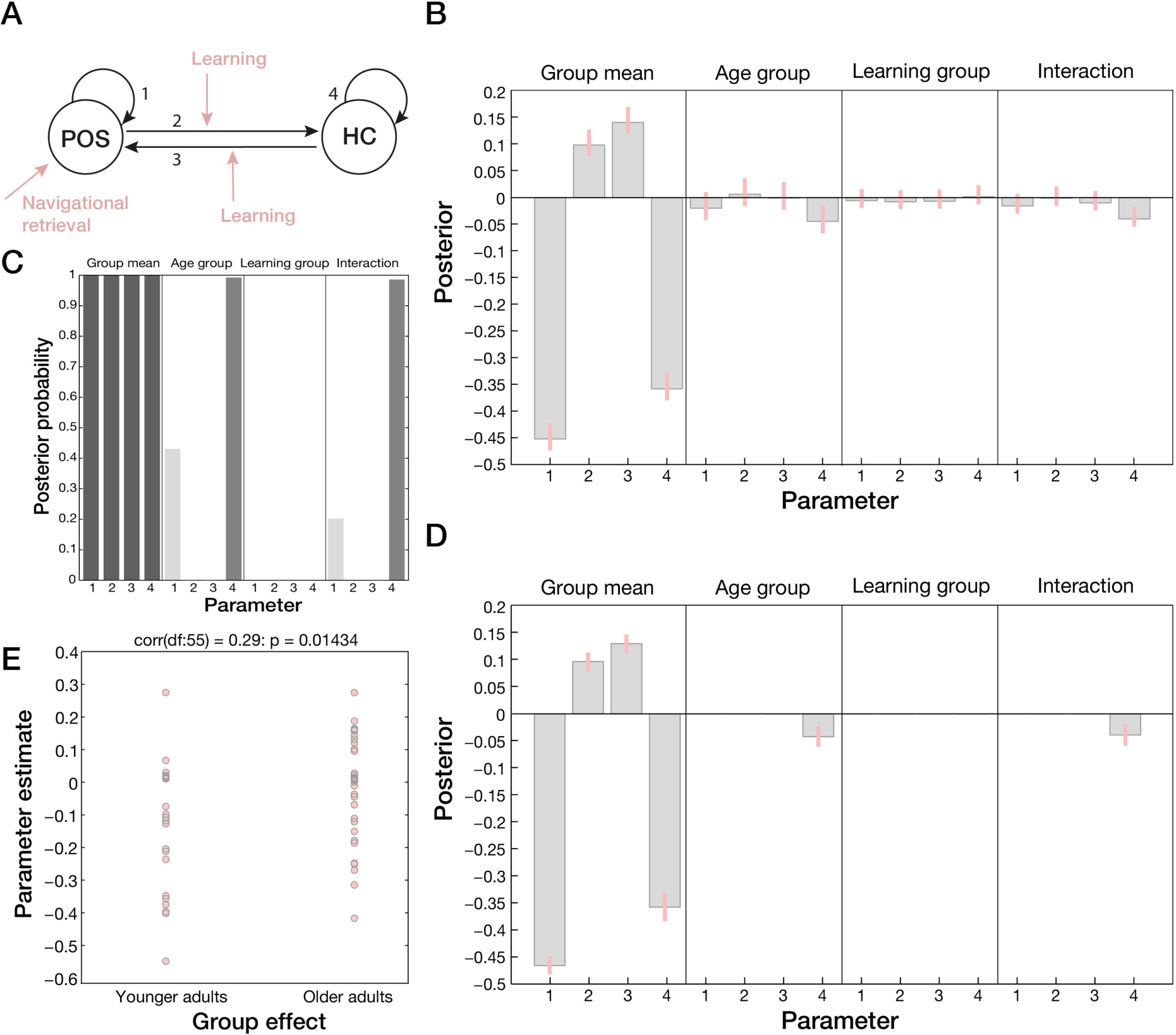
Results of the DCM PEB analysis. (A) First-level DCM specification to determine average connectivity within and between anterior hippocampus and POS. Navigational retrieval phases were modeled as driving input entering the cortical network via the POS, and the amount of learning per block was included as modulatory input on the bidirectional connections between the regions. Estimated Parameters (1: self-connection POS, 2: POS – hippocampus connection, 3: hippocampus – POS connection, 4: self-connection hippocampus) (B) before and (D) after Bayesian model reduction (BMR) for each covariate (age group, learning group, interaction between age group and learning group) in the second-level PEB model. Grey bars represent parameter means and pink lines their 95% confidence intervals. The parameters for self-connections (parameter 1 and 4) are expressed as log scaling parameters that can be converted to Hz using x_Hz = −0.5 * exp(x) whereby x is the log scaling parameter and −0.5 Hz the prior. (C) Posterior probabilities per parameter for each second-level covariate after BMR, (E) Predicted age group of each participant as derived from a LOO cross-validation scheme based on the estimated self-connection strength in the anterior hippocampus.

With respect to age group differences in connectivity, only one parameter survived BMR (second panels of Figure 6B and 6D). Specifically, older compared to younger adults had a reduced inhibitory self-connection strength in the anterior hippocampus, i.e., a relative disinhibition in this region. Note that for self-connections in the DCM framework, parameters are expressed as log scaling parameters and that the regressor representing age group was coded in a way that the resulting parameter is the amount that needs to be added to the group mean to obtain the older adults’ connection strength (the group mean is obtained by calculating −0.5Hz * exp(−0.33698) = −0.357Hz and for older adults −0.5Hz * exp(−0.33698 + −0.039719) = −0.3431Hz). Thus, our model provides evidence that the aging hippocampus seems to be more readily excited by afferent activity from other regions during spatial learning. The interaction between age group and learning sub-group in this model parameter also survived BMR (right panels in Figure 6B and 6D). Inspection of the fourth panel in Figure 6B indicates that the modulation of the hippocampal self-connection strength due to aging was stronger than its modulation due to overall learning ability of the individual (see also Figure 6C for posterior probabilities of each parameter). We did not find any modulatory effects of the (within-subject) amount of learning per block.

We further performed a leave-one-out (LOO) cross-validation using the model parameter denoting the self-connection strength in the anterior hippocampus to test whether this effect would be large enough to predict the participants’ age group. In this analysis, all but one subject are used to estimate the model parameter, which is then used to evaluate the posterior belief of the model parameter in a left-out (test) subject. The predicted and actual between-subject effect for each test subject is then compared to derive an independent out-of-sample correlation, which was 0.29 in the current sample (p=0.01434, Figure 6E). Thus, the estimated intrinsic connection strength in the anterior hippocampus during spatial learning was large enough to predict the age group of a new subject above chance level.

## DISCUSSION

In this study, we show that healthy older adults, on average, have substantial problems in learning to orient themselves in a novel, city-like virtual environment. In addition, we found that their ability to retrieve the spatial layout of the environment is more rigidly tied to specific viewpoints encountered at the beginning of learning compared to younger adults. We show that the individual’s mean amount of learning and the increase in reaction times constitute two distinct behavioral features that can be used to predict the age group of the individual during spatial learning. At neural levels, we could replicate earlier findings in younger adults showing that activity in RSC/POS increases while activity in the anterior hippocampus decreases as a function of learning (Auger et al., 2015; Brodt et al., 2016; Wolbers & Büchel, 2005). We provide novel evidence that in older adults, activity in these two regions is decoupled from the amount of learning and does not change systematically across repeated episodes in the environment. Importantly, we provide the first evidence that an increased excitability of the anterior hippocampus, as demonstrated by an age-related reduction of its inhibitory self-connection strength, might constitute a potential neural mechanism for these changes.

### Age-related differences in retrieving spatial information during early learning stages

The observed performance differences between our age groups are in line with findings from previous behavioral studies (Iaria et al., 2009; Yamamoto & DeGirolamo, 2012). By applying a standard health screening procedure and broadly assessing cognitive performance, we ensured that our final samples did not show any evidence for major cognitive impairment. Thus, spatial learning is a cognitive skill that is subject to substantial decline in the majority of older adults, even if they are otherwise healthy. However, we cannot rule out that some of the older adults may already exhibit preclinical signs for AD that might have affected their performance. For example, an increased Aβ deposition or tau pathology are known to occur in brain regions that are involved in spatial navigation during the earliest phase of AD (Braak & Del Tredici, 2015).

Importantly, by changing the directions from which locations were approached from the first to the second half of the experiment, we could show that older adults are less flexible to account for slight changes in sensory input during retrieval. This might be related to age-related deficits in distinguishing novel from familiar input and the successful retrieval of already stored information that has been linked to computations in the hippocampus (i.e., pattern separation and pattern completion; Vieweg, Stangl, Howard, & Wolbers, 2015; Yassa et al., 2011). Alternatively, age-related deficits in integrating different viewpoints or in allocentric processing more generally, as shown in Wiener, de Condappa, Harris, and Wolbers (2013), might have contributed to this pattern of results. One should note, however, that this higher reliance on the sensory input during retrieval became apparent in the form of a surprise signal (i.e., longer response times after the change in directions took place) but did not affect task accuracy. This is in contrast to the findings reported in Wiener et al. (2013) who found that performance in older adults is impoverished when locations are approached from novel directions during route learning. The authors suggested that this might be related to the adoption of a response strategy in which landmarks are used as cues that become linked to a specific motor response. In the current study, we did not find indications for any biases in pointing behavior (i.e., a preference for pointing along streets) in older adults, which would suggest that they might have tried to use a response strategy. Previous research showed that viewpoint independence in spatial memory is supported by the hippocampus (King, Burgess, Hartley, Vargha-Khadem, & O’Keefe, 2002). Thus, our behavioral results already point to an impaired information processing within the hippocampus such that older adults had more problems in handling shifted viewpoints and storing a viewpoint independent spatial representation of the environment.

### Impaired information processing in the aging anterior hippocampus and POS/RSC

Our replication of findings from previous studies concerning learning-related activity changes in the RSC/POS and the hippocampus of younger adults (Auger et al., 2015; Brodt et al., 2016; Wolbers & Büchel, 2005) shows that our task was suitable to measure spatial learning, while using a complex photorealistic VE. In older adults, activity in these regions was generally increased compared to younger adults when contrasted to our control condition and did not change systemically as a function of learning over the course of the experiment. Thus, one factor that determines whether activity decreases or increases are observed in older versus younger adults relates to the participant’s learning status at the time-point when activity is measured. This might be one explanation why our results are in contrast to previous findings in humans who observed activity decreases in the RSC and hippocampus during spatial navigation (Moffat et al., 2006). Moreover, Moffat et al. (2006) measured brain activity during encoding when participants were actively navigating in VR, and they controlled for performance differences in their age-group comparisons. In the present study, we focused on neural activation patterns during retrieval phases that followed encoding phases to measure the amount of learning that was acquired in each encoding phase. Thus, another factor that might have contributed to our findings might be the overall task demands given that we also observed an age-related activity increase in RSC/POS and hippocampus when contrasting retrieval phases to encoding phases.

We further found age-related activity reductions in superior and lateral parts of the POS when comparing navigation versus control. The anatomical definition of the RSC varies between studies, and sometimes RSC proper (Brodmann areas 29 and 30) and POS are grouped together and are collectivity referred to as retrospenial complex (Epstein, 2008; Vass & Epstein, 2013). However, each region’s role during spatial navigation might differ. Previous research indicates that RSC is primarily involved in the processing of landmark permanence, whereas POS may relate landmarks to certain locations and might show a different time-course of its involvement during spatial learning (Auger et al., 2015). In the present study, we found a learning-related activity increase in RSC proper as well as in clusters in the POS, located very closely to previously reported activations (Brodt et al., 2016; Wolbers & Büchel, 2005). Disentangling the respective role of different parts of RSC and POS during spatial navigation in the aging brain would be an important topic for future research.

Notably, an age-related hyperactivity in the hippocampus has been observed in several studies investigating age-related deficits in pattern separation in humans (Reagh et al., 2018; Yassa et al., 2011), as well as in rodent and non-human primate studies on age-related changes in spatial navigation (Thomé et al., 2016; Wilson et al., 2005). By examining age-related changes in effective connectivity, we were able to show, for the first time, that a reduction in the inhibitory self-connection strength of the anterior hippocampus (i.e., a relative disinhibition) might constitute the underlying neural mechanism for the elevated signal in this region. This indicates that the aging hippocampus is more sensitive to input from other regions during spatial learning, which is in line with findings that increased firing rates in CA3 place cells of memory impaired monkeys is linked to a reduced number of GABAergic inhibitory interneurons that regulate the overall excitability of the network (Thomé et al., 2016). This in turn might lead to higher firing rates and errors in distinguishing between similar spatial representations during retrieval.

Within the context of DCM, the self-connection in a brain region has been conceptualized as modulating the gain or excitability of neuronal populations reporting mismatches between signals from different levels of the cortical hierarchy (i.e., prediction errors), reflecting the precision of the prediction errors (Clark, 2013; Friston et al., 2017). Thus, an increased excitability of the anterior hippocampus does not seem to constitute a compensatory mechanism in the aging brain. In line with this, we found evidence that this age effect outweighs the effects of learning on the self-connection strength in the anterior hippocampus. Bakker, Albert, Krauss, Speck, and Gallagher (2015) showed that hyperactivity in the dentate gyrus and CA3 subfield of amnestic patients can be reduced by applying low doses of an anti-epileptic drug that targets excitatory neurotransmission, which in turn improves memory performance. In aged rats, it has similarly been shown that treatments to reduce hippocampal hyperactivity improve memory performance and spatial information processing (Koh, Haberman, Foti, McCown, & Gallagher, 2010; Koh, Rosenzweig-Lipson, & Gallagher, 2013; Robitsek, Ratner, Stewart, Eichenbaum, & Farb, 2015). The increased excitability of the aging hippocampus may consequently also impair information processing in RSC/POS during spatial learning, given the interactions between the two regions. In line with this, Mao et al. (2018) found that bilateral hippocampal lesions impair place cell activity in RSC and suppress the gradual emergence of a spatial code in this region.

### Inter-individual differences in learning within the older age group

Performance in our task was highly variable between individuals. More specifically, while some older adults learned the layout of the environment as quickly as younger adults, we found that others showed continuous learning across the whole experiment, learned very slowly, or were not able to retrieve relevant information to perform the task. Because the aim of our study was to measure the process of learning a novel spatial environment during fMRI scanning, we kept the amount of exposure in the VE constant between the age groups while ensuring that overall scanning time was still reasonable. Therefore, we cannot determine whether older adults would just need more time for learning. However, in the light of the large heterogeneity in performance within the older age group, it seems unlikely that all of them would have reached the same level of performance as the younger adults if provided with more time in the VE. Using machine learning methods on MRI data of hundreds of older adults, Eavani et al. (2018) recently described multiple phenotypes of resilient and advanced brain agers that are characterized by specific functional and structural changes. Interestingly, the authors described one phenotype that displays atrophy in the hippocampus, decreased coherence in posterior parts of the medial parietal cortex, and an increased connectivity in the MTL. Thus, older adults who show an increased excitability of the anterior hippocampus might be particularly impaired in memorizing novel spatial environments, even if they spend a lot of time exploring them. This increased excitability may further constitute a precursor for subsequent changes at neural levels that are known to advance the progression to AD (cf., Leal et al., 2017).

Finally, by using the estimated hidden learning states of the participants to form sub-groups of learners in our sample and including this information in the analysis of the fMRI data, we found that within older adults, activity in the posterior hippocampus differed as a function of the overall learning ability of the individual. In contrast to those groups who showed less learning across the experiment, this portion of the hippocampus was more involved in those older adults who performed at ceiling and were able to generate a spatial representation of the environment quickly. It is well known that anterior and posterior hippocampus represent information at different scales with representations in the anterior hippocampus being rather coarse, while information in the posterior hippocampus is represented in more fine-grained detail (Poppenk, Evensmoen, Moscovitch, & Nadel, 2013). This also relates to the familiarity of the processed input with information that is more familiar being represented in the posterior hippocampus, whereas novel information is typically processed in anterior hippocampus at a lower resolution. In line with this, it has been shown that high levels of expertise in spatial navigation are associated with greater volumes in the posterior hippocampus (Maguire et al., 2000). Moreover, APOEε4 status has been associated with changes in functional connectivity that are particularly evident in anterior parts of the hippocampus in older adults during memory performance (Harrison, Burggren, Small, & Bookheimer, 2016). Thus, one might speculate that in some older adults, recruitment of the posterior hippocampus might be able to compensate for compromised functioning of the anterior hippocampus, which is associated with navigational performance that is comparable to that of high-performing younger adults. Future studies should examine the causes of this high heterogeneity in older adults in more detail and consider the contributions from different hippocampal subfields and apply additional measures, for example preclinical markers for AD, to further characterize the neural mechanisms for age-related deficits in spatial learning and, specifically, why these abilities are preserved in some older adults.

### Conclusion

Increased excitability of the anterior hippocampus, which may also affect RSC/POS functioning, provides a novel explanation why older adults have problems to form an accurate spatial representation of a newly encountered environment. In addition, our findings add to a growing body of evidence associating hyperactivity in the hippocampus to memory impairments in older humans, non-human primates, and rodents and link it to preclinical markers for AD. Our findings have important implications for future research investigating the potential of individual learning states during spatial navigation as a cognitive marker for the progression from healthy aging to neurodegenerative disease.

## MATERIAL AND METHODS

### Participants

In the behavioral experiment, 17 younger (9 female, mean age: 24.0 ± 1.66, age range: 21-28) and 17 older adults took part (8 female, mean age: 66.4 ± 2.69, age range: 61-72). All of them were right-handed (LQ: 91.9 ± 11.0; Oldfield, 1971) and the older adults showed no signs of major cognitive impairment with scores higher than 23 in the Montreal Cognitive Assessment (MoCA score: 26.9 ± 2.18; Luis, Keegan, & Mullan, 2009; Nasreddine et al., 2005).

In the fMRI experiment, a total of 64 participants (27 younger adults, 37 older adults) took part. Three participants (one younger and two older adults) were excluded from further analyses because they were identified as outliers in the fMRI data quality checks. In addition, one younger and three older adults were excluded due to problems in following task instructions and/or cybersickness. The final fMRI sample consisted of 25 younger (13 female, mean age: 23.4 ± 2.18, age range: 20-26) and 32 older adults (17 female, mean age: 67.3 ± 4.80, age range: 58-75). They were all right-handed (LQ: 90.4 ± 12.1; Oldfield, 1971) and the older adults did not show signs of major cognitive impairment (MoCA score: 27.6 ± 1.93, range: 25-31; Nasreddine et al., 2005).

Across experiments, participants had normal or corrected-to-normal vision and none of them reported a history of psychiatric or neurological diseases or use of medication that might affect task performance or fMRI scanning. In addition, most of the participants already participated in previous virtual reality (VR) experiments at the DZNE and, hence, were familiar with navigating in these kinds of setups. Participants provided informed consent and were paid for their participation in accordance with the local ethics committee.

### Virtual Environment

Using 3ds Max (Autodesk, San Rafael, CA, USA), a novel virtual environment (VE) was developed, which resembled a typical German historic city center consisting of town houses, shops and restaurants. The VE had a square-like spatial layout with four interconnected 4-way intersections (Figure 1B). At two intersections, a church and a town hall were placed at the end of one of the outgoing streets, whereas a 2D wall displaying a photo texture of a street continuation bordered the remaining street ends. The VE was based on a 3D model of the old city center of Tübingen, which lies approximately 450 km south of Magdeburg where data collection took place. All of the participants confirmed to have never visited Tübingen before the time of testing.

### Behavioral Task and Procedure

Vizard 5.0 (World Viz, Santa Barbara, CA, USA) was used to animate the experiments, which both started with a familiarization phase during which the participants encountered the VE for the first time. Their task during this phase was to collect tokens that were placed at the street ends by moving around the VE, using the four arrow keys of a standard computer keyboard. This phase ended once every token was collected, ensuring that they had visited every street at least once. It followed a short practice of the pointing task (8 trials) that was used to measure navigational retrieval in the experiments. In this way, the VE and the task were introduced in a step-wise manner to reduce the impact of different degrees of experience in handling VR setups on task performance (Diersch & Wolbers, 2019).

In the behavioral experiment, eight learning blocks were implemented during which eight retrieval phases alternated with seven encoding phases. One navigational retrieval phase consisted of 12 pointing trials. A pointing trial started with participants being passively transported towards one of the intersections (I1-I4 in Figure 1B) starting from one of the four streets leading towards that intersection (D1-D4 in Figure 1B). Duration of this travel phase was fixed to 4 s corresponding to 20 virtual meters. The movement stopped at the center of the intersection, a red crosshair appeared in the middle of the screen, and participants were asked to point in the direction of one of the two target landmarks (see Figure 1C and Video 2 for an example trial). Pointing was performed by moving the crosshair to the left or right with the arrow keys of the keyboard. Once they believed to have reached the correct position, they confirmed their response by pressing the space bar. Participants were asked to respond as fast and accurately as possible with a time-out of 12 s (corresponding to 1½ 360° turns in the VE). The ITI, showing a fixation cross, was fixed to 1.5 s. Throughout each trial, a picture cue of the target landmark was displayed at the bottom of the screen, and the background was obscured by fog to prevent participants from seeing the street ends or target landmarks during pointing. The first seven retrieval phases were followed by an encoding phase during which participants were passively transported around the whole VE (without fog), starting from one of the two target landmarks in clockwise or counterclockwise order, counterbalanced across the experiment (see Figure 1B for one example encoding tour and Video 1 for a short segment of one tour). During encoding, participants were instructed to pay close attention to the spatial layout of the VE and the location of the target landmarks. Passive transportation instead of self-controlled traveling was chosen to keep the duration of the encoding tours constant for both age groups. In total, participants performed 96 navigational retrieval trials (4 intersections × 4 directions × 2 target landmarks × 3 repetitions) in a pseudo-randomized order, with the restriction that each intersection/target landmark combination was encountered starting from two of the four possible directions in the first half of the experiment. In the second half of the experiment, divided by a self-timed break, the respective other two directions were used, counterbalanced across participants.

The fMRI experiment also consisted of eight learning blocks during which eight retrieval phases alternated with seven encoding phases (Figure 1A). fMRI scanning started after a familiarization phase outside of the scanner with the same structure as in the behavioral experiment and a short practice phase during structural imaging. One retrieval phase consisted of 8 navigational retrieval trials, which were followed by 4 control trials. These control trials also started with a 4 s travel phase towards an intersection, followed by a pointing phase with a crosshair on screen. Here, cued by a corresponding picture, however, participants were instructed to indicate which of the four corner buildings at the intersection had changed its color and was shaded in blue. Their responses in the control task were classified as correct if they lied within ± 25° from the middle of the respective building, approximately corresponding to its outline. Participants moved the crosshair with their index and middle finger for left and right turns and confirmed their responses with their thumb on a 5-key Lumitouch response box that was placed in their right hand during scanning. Again, participants were asked to respond as fast and accurately as possible with a time-out of 12 s. The ITIs had a variable duration of 1-5 s with a mean of 3 s. During retrieval trials, an additional jittered interval of 0.5-1.5 s duration with a mean of 1 s was included after the travel phase/before the crosshair appeared. The structure of the respective encoding tours was the same as in the behavioral experiment. In total, participants performed 64 navigational retrieval trials (4 intersections × 4 directions × 2 target landmarks × 2 repetitions) without the change of directions from the first to the second half of the experiment as in the behavioral experiment. They additionally performed 32 control trials (4 intersections × 4 directions × 2 repetitions). fMRI scanning consisted of 3 runs that were divided by short breaks with 24 navigational retrieval trials, 12 control trials and 2 encoding tours in the 1^st^ run; 24 navigational retrieval trials, 12 control trials and 3 encoding tours in the 2^nd^ run; and 16 navigational retrieval trials, 8 control trials and 2 encoding tours in the 3^rd^ run.

### Bayesian Modeling of Performance Data

Subject-specific improvements in navigational performance were estimated by using a Bayesian implementation of a state-space model that is similar to a local level model where the trial outcomes, y, correspond to the observed level, and the state level represents the hidden learning state, μ (Figure S1; Commandeur & Koopman, 2007). The hidden learning state, μ, is following a random walk such that the actual block learning state depends on the learning state from the previous block. Similar state-space models (e.g., Smith, Wirth, Suzuki, & Brown, 2007) have been used in previous studies to estimate spatial learning (Auger et al., 2015; Wolbers & Büchel, 2005). However, these studies modeled binary data on a trial-by-trial basis, whereas the present study used continuous performance outcomes and focused on estimating spatial learning block-wise instead of trial-wise. To model the learning state block-wise, an intermediate level accounts for the effects of the responses, η, and shrinks the effects of individual trials within a block towards the block-wise learning state. In this way, the model accounts for the fact that we can only measure behavioral performance but not the effect of learning or navigational improvement, which we expected to change from one encoding phase to the next but not necessarily from trial to trial. Introducing this intermediate level additionally allowed us to incorporate potential missing trials into the response effects, η. In case of missing trials, we estimated η ∼ HalfNormal 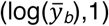, i.e., using the log of the block mean as location parameter. The model was implemented using the Python interface to Stan, PyStan (Carpenter et al., 2017; Stan Development Team, 2017; see Figure S2 for the Stan code). To account for the substantial between-and within subject variability of the data, weakly informative priors were chosen to provide vague guidance for effective sampling. The model was fit for each participant using four chains each with 4000 iterations, of which 2000 correspond to the warm up period, totaling 8000 post-warm-up draws. After inference, convergence of the chains was checked by means of the effective sample size and the potential scale reduction factor (Rhat), confirming that our chains mixed well (Gelman & Shirley, 2011).

To determine the fit of our model to the data, we performed a posterior predictive check that compares the observed data with simulated data using samples from the posterior distribution. In Figure S3A-E, the posterior predictive samples distribution y_rep_ is plotted together with the observed data y for representative individuals from different learning sub-groups (see <u>Performance Clustering</u> below) showing that our model was adequate to capture the observed data. We further compared our model to an alternative, simpler model where η was removed (i.e., learning was estimated trial-wise instead of block-wise). Using a leave-one-out (LOO) cross-validation (Vehtari, Gelman, & Gabry, 2017), point-wise out-of-sample prediction accuracies were estimated for both models. Comparing them confirmed that the model incorporating the intermediate layer accounting for the response effects, η, provided better fit to the data, as evidenced by positive LOO differences across the whole sample (sample mean = 1209, SE = 242; see S3F for a histogram showing the individual LOO difference values).

### fMRI Acquisition Parameters

Scanning was performed on a 3T Magnetom Prisma scanner (Siemens Healthcare, Erlangen, Germany) with a 20-channel head coil. High-resolution T1-weighted anatomical images were acquired using a MPRAGE sequence (1 mm isotropic resolution; TE = 2,82 ms; TR = 2500 ms; flip angle = 7°). In three functional runs, whole-brain T2*-weighted echo planar images with BOLD contrast were acquired in interleaved bottom-up order (36 slices, 3 mm isotropic resolution; TE = 30 ms; TR = 2000 ms; FoV = 216 mm; 72 × 72 image matrix; flip angle = 90°).

### Behavioral and fMRI analyses

#### Behavioral Analyses

Absolute pointing errors (i.e., the deviation of the subject’s response from the respective target landmark) served as performance measures in both experiments. In the behavioral experiment, we additionally analyzed response times given the change in directions from which the intersections were approached after the first half of the experiment. Where appropriate, analyses of variance (ANOVA) were performed across learning blocks with age-group (younger adults, older adults) as between-subjects’ variable. Circular statistics were applied on the signed pointing error data relative to each target landmark for every intersection-direction combination, using the CircStat toolbox in MATLAB (Berens, 2009). In general, a threshold of p < 0.05 was considered significant (with correction for the number of tests where applicable).

#### Logistic Regression Model

With respect to the behavioral experiment, we were interested in whether two features that characterized age-related differences in performance could be used to predict the age group of our participants. The first feature was the mean amount of learning across all learning blocks, which was calculated based on the differences between individual learning state estimates, derived from the Bayesian state-space model, from two consecutive learning blocks. The estimates from the first learning block after the familiarization phase, during which participants encountered the VE for the first time, were subtracted from chance level performance (90°). In this way, learning related improvements in performance were considered that already took place during the familiarization phase, resulting in pointing errors well below chance level in the first learning block for some participants. The second feature were the changes in response times after the directions changed from which the intersections were approached after the first half of the experiment. These two features were consequently normalized and then fed into a logistic regression model as implemented in Scikit-learn (Pedregosa et al., 2011), with age group as target vector. The regularization parameter was set using a 10-fold nested cross-validation, and the performance of the model was assessed by computing the average area under the curve (AUC) for all folds. In this way, the probability of each individual actually belonging to the younger or the older age group could be estimated. The resulting probabilities are interpreted in terms of individual performance: those participants with a higher probability of belonging to the younger age group show better performance on the task while a higher probability of being in the older age group relates to poorer navigational performance.

#### Performance Clustering

In the analysis of the behavioral data from the fMRI experiment, we assessed whether subjects could be clustered into different learning sub-groups based on their performance. This allowed us to investigate learning-related differences in neural activity at the between-subjects level. For each participant, we created a distribution based on the difference of the latent state distributions of the last and first learning block to capture the overall amount of learning across the experiment. The mean and the standard deviation parameters of this difference distribution were obtained by fitting it to a normal distribution using SciPy (Jones, Oliphant, & Peterson, 2001). We used a K-means clustering algorithm as implemented in Scikit-learn (Pedregosa et al., 2011) to identify the centers of a pre-determined number of clusters based on their distances to the data points. To obtain the optimal number of clusters to input into the K-means, we computed the silhouette score varying the number of possible clusters from 3 to 7 and found 5 to be the best choice for the number of learning sub-groups in our sample.

#### fMRI Image Quality Control and Preprocessing

The imaging data were first transformed into the Brain Imaging Data Structure (BIDS) format (Gorgolewski et al., 2016). MRIQC (Version 0.9.3; Esteban et al., 2017) was used for checking the quality of the MRI data. MRIQC utilizes tools from different software packages such as FSL or ANTs to extract image quality metrics (IQMs) and generates visual reports at the individual and group level. This allows the evaluation of different characteristics of the structural and functional MR images, for example, SNR/tSNR, sharpness, and presence of artifacts. Data from one younger adult and two older adults were consequently excluded from further analyses due to strong task-related movement and/or artifacts in several functional runs resulting in low-quality IQMs (e.g., high Ghost-to-Signal ratio, low tSNR). Next, preprocessing was performed using fMRIprep version 1.0.0-rc5 (Esteban et al., 2019) that also draws on software packages such as FSL, ANTs, AFNI, and FreeSurfer to provide the optimal implementation for different stages of preprocessing, including motion correction, slice timing, co-registration, and normalization. Finally, the data were smoothed with a 6 mm full-width at half maximum isotropic Gaussian kernel using SPM 12 (Wellcome Department of Imaging Neuroscience, London, UK).

#### ROI Definition

Based on results from previous studies (Auger et al., 2015; Mao et al., 2018; Wolbers & Büchel, 2005), we defined two regions of interest (ROI), namely, the RSC/POS and the hippocampus. The single ROIs were created based on each participant’s T1 structural scan using a semiautomated anatomic reconstruction and labeling procedure as implemented in FreeSurfer v6.0.0, which is part of the fMRIprep pipeline (http://surfer.nmr.mgh.harvard.edu; Dale, Fischl, & Sereno, 1999; Fischl, Sereno, & Dale, 1999). In each hemisphere, labels corresponding to the posterior-ventral part of the cingulate gyrus (area 10) and the parieto-occipital sulcus (area 65) from the Destrieux Atlas and the hippocampus from the subcortical segmentation were extracted (Destrieux, Fischl, Dale, & Halgren, 2010; Fischl et al., 2002). The two cortical labels were combined into one RSC/POS ROI. Each ROI was next transformed to MNI space. Hemispheres were combined to one bilateral ROI, thresholded at 0.5, and finally resampled to correspond to the resolution of our functional images. The ROIs were subsequently used in the univariate analysis and for the volumes of interest (VOI) extraction in the effective connectivity analysis (see below).

#### fMRI Univariate Analysis

At the single-subject level, a general linear model (GLM) was specified with six regressors of interest for each learning block using a high-pass filter of 100 Hz. For the navigational as well as control retrieval trials, we created regressors for the 4 sec travel phase and the pointing phase. For the encoding phases, regressors modeled periods when participants were located within 20 m of the intersection centers (corresponding to the area covered during the retrieval travel phases) as well as outside of these areas. Finally, the time of the button press was modeled as regressor of no interest. All regressors were convolved with the standard canonical hemodynamic response function (HRF) in SPM12. In addition, we included motion parameters, the frame-wise displacement (FD) and physiological noise regressors (aCompCor values; Behzadi, Restom, Liau, & Liu, 2007), as obtained from fMRIprep preprocessing, in the GLM to control for physiological and movement confounds. We first contrasted navigational retrieval trials to control trials across learning blocks to identify general activation patterns in the RSC/POS and the hippocampus during spatial navigation in our complex real-world environment, similar to previous studies investigating age-group differences in spatial navigation (Moffat et al., 2006). We additionally contrasted the travel phases towards the intersections during navigational retrieval trials to the corresponding periods when participants encountered the same areas during the encoding tours. The within-subject effects of learning were assessed by using the normalized differences between learning state estimates from consecutive learning blocks (i.e., amount of learning) as contrast weights over the regressors modeling each travel phase during navigational retrieval per learning block (cf., Wolbers & Büchel, 2005). At the group level, the resulting individual contrast images were entered into two-sample t-tests to assess interactions with age group. Finally, in order to check in which regions activity changes across learning blocks are modulated by the overall learning ability of the subject, we ran an additional analysis in which learning sub-group was added as covariate. We focus on activations that survived a FWE-correction for multiple comparisons using a threshold of p < 0.05 at the cluster level.

#### Effective Connectivity Analysis

Effective connectivity within and between the hippocampus and the POS was examined using the parametric empirical Bayesian (PEB) approach in the context of Dynamic Causal Modeling (DCM) as implemented in SPM12 (Friston et al., 2016).

#### GLM and VOI Selection

For the DCM analysis, we created a second GLM in which the time-series from our three functional runs were concatenated and added regressors that modeled the mean signal for each run. The amount of learning per learning block was included as parametric modulation of the regressor modeling the travel phase during navigational retrieval trials for each participant. All other regressors were the same as in the first GLM although they were not modeled separately for each learning block. The sanity check of the concatenated GLM revealed that activity in the right anterior hippocampus (27, −9, −15, Z = 3.97; 27 voxels) decreased and activity in the left POS (−12, −63, 31, Z = 3.56; 42 voxels) increased with the amount of learning in younger adults (p < 0.05, FWE-corrected for the respective ROI). No additional activations emerged elsewhere in the brain. When testing for interactions between learning-related activity changes and age group within our ROIs, one cluster within bilateral POS extending to RSC was revealed (15, −66, 44, Z = 4.23; 12, −57, 4, Z = 4.21; −6, −66, 24, Z = 3.94; −15, −60, 28, Z = 3.61; 369 voxels). Thus, activity in this region increased with learning in younger adults but less so in older adults. The slight differences of these results to the ones from the first GLM are likely related to differences in the design of the two GLMs. Whereas the first GLM was optimized to capture our experimental design as precisely as possible by modeling all regressors of interest separately for each learning block, the concatenated GLM was optimized for the DCM analysis that relies on single-run time-series.

BOLD time-series were extracted for each individual using a t-contrast over the regressors modeling the travel phase during navigational retrieval and the amount of learning with a liberal threshold of p < 0.1 (Note that this threshold was only used for VOI selection, but not in the final DCM statistics). The principal eigenvariate was extracted around the group peak coordinates within the hippocampus and POS as obtained in the univariate analysis of the concatenated GLM and was allowed to vary as a 8 mm sphere centered on the subject-specific maximum constrained by a 24 mm sphere centered on the group maximum and the respective ROI mask. In this way, variation between individuals in the exact location of the effect was taken into account, given the high heterogeneity in our sample and slightly different peak voxels in the two GLMs. The extractions were corrected using an F-contrast that retained the effects of interest (navigational as well as control retrieval phases, encoding phases, button press) while partitioning out task-unrelated variance caused by head motion, for example. For participants for which no supra-threshold voxels were identified (three younger adults and one older adult), the threshold was dropped to p < 0.5 to extract BOLD time-series (cf., Zeidman, Jafarian, Corbin, et al., 2019).

#### First-Level DCM Specification

We specified a bilinear, one-state DCM for each participant by setting the regressor modeling the travel phase during navigational retrieval trials as driving input entering the cortical network via the POS. The amount of learning per learning block was included as modulatory input on the bidirectional connections between hippocampus and POS (Figure 6A). All inputs were mean-centered so that the A-matrix of the DCM represents the average connectivity across experimental conditions. We used stochastic DCM that seeks to improve model estimation by modeling random fluctuations and hidden neuronal causes in the differential equations of the neuronal states (Daunizeau, Stephan, & Friston, 2012; Li et al., 2011). In this way, the impact of potential confounding effects of variations in BOLD response caused by age is reduced. Bayesian group inversion was performed, providing estimates of the connection strength parameters that best explained the observed data per participant. Critically, within DCM PEB, at each iteration of the within subject inversion, the individual priors are updated using the group average connection strengths as priors. Inspection of the single DCMs after inversion confirmed that our full model provided good fit to the observed data with an average of 44.5 ± 3.22% variance explained.

#### Second-Level PEB Model

Next, we created a second level PEB model over the parameters that included the group mean and age group as covariates to identify differences between younger and older adults. We further included learning sub-group and its interaction with age group as covariates in the model. To calculate the interaction term, the regressors for age group and learning sub-group were mean-centered. A search over nested PEB models was performed by using Bayesian model comparison (BMC) that explores a space of models under the assumption that different combinations of the connections may exist across participants (Zeidman, Jafarian, Seghier, et al., 2019). To search over hundreds of nested models incorporating different combinations of connections and group differences, BMR is used that iteratively prunes parameters from the full model until model-evidence decreases. To reduce dilution of evidence, we separately checked for group differences in the A-matrix (average connectivity across experimental conditions) and the B-matrix (within-subject modulatory input of the amount of learning per block). We further performed a LOO cross-validation to check whether the model parameter that differed between older and younger adults could be used to predict the participants’ age group.

#### Data Availability

Source data files for the main results figures and tables are stored at https://osf.io/fjbxu/. We additionally provide a key resources table listing all the software packages that were used in the current study. The Stan code of the Bayesian state-space model can be found in Figure S2.

## ACKNOWLEDGMENTS

We thank Tobias Meilinger from the MPI for Biological Cybernetics, Tübingen, Germany for providing us with their 3D model of the Tübingen city center that served as basis for our virtual environment, Peter Zeidman for his help with the DCM PEB analysis, Johannes Achtzehn for his help with 3D modeling and programming of the experiments, Lena Wattenberg, Judith Siegel, and Katharina Mamsch for assistance in data collection.

## AUTHOR CONTRIBUTIONS

Conceptualization, N.D. and T.W.; Project administration, N.D.; Investigation, N.D.; Resources, T.W. and C.T.; Methodology & Formal Analysis, N.D. and J.P.V.-H.; Visualization, N.D.; Writing – Original Draft, N.D.; Writing – Review & Editing, N.D., J.P.V.-H., C.T., and T.W.; Supervision, T.W.; Funding Acquisition, T.W.

