## Supplementary Material for "Increased hippocampal excitability and altered learning dynamics mediate cognitive mapping deficits in human aging"

### SUPPLEMENTAL INFORMATION

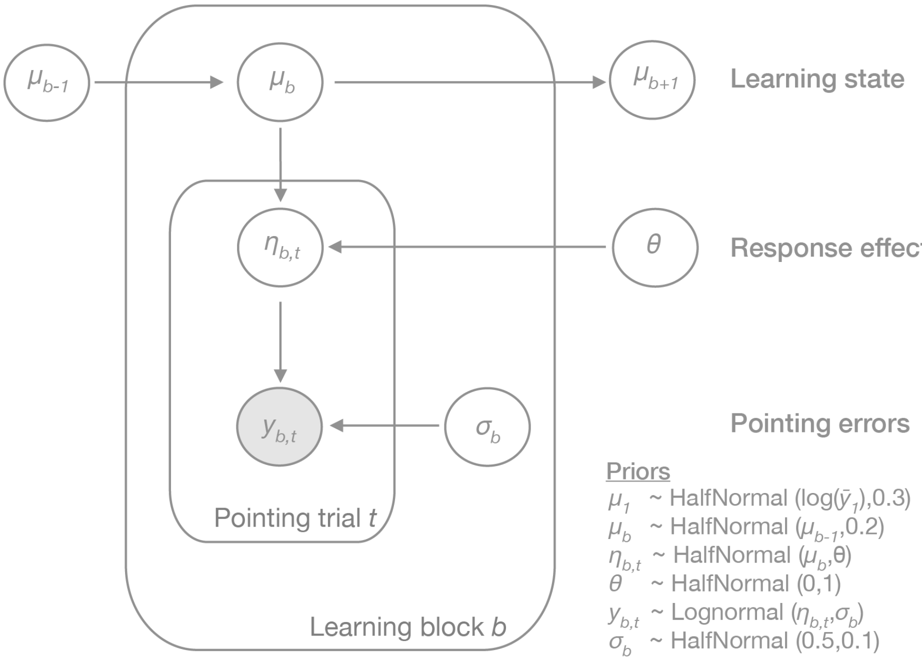


**Figure S1, related to Figure 2, Figure 3, and Bayesian Modeling of Performance Data section:** Bayesian state-space model to estimate the subject-specific hidden learning state per learning block.

```
data {
  int<lower=0> nblocks;
  int<lower=0> ntrials;
  matrix[nblocks, ntrials] y;       // the data
  int<lower=0, upper=1> fit_model;  // whether to fit data or not
  // vars controlling the priors
  real theta_loc;
  real theta_scale;
  real mulog1_loc;
  real mulog1_scale;
  real mulog_scale;
  real sigma_loc;
  real sigma_scale;
}

parameters {
  real<lower=0> theta; // theta is also *regularizing* the random effects
  vector<lower=0,upper=log(180)>[nblocks] mulog;
  matrix<lower=0>[nblocks, ntrials] eta;
  vector<lower=0>[nblocks] sigma;
}

model {
  theta ~ normal(theta_loc, theta_scale);
  mulog[1] ~ normal(mulog1_loc, mulog1_scale);
  for (b in 2:nblocks) {
    mulog[b] ~ normal(mulog[b - 1], mulog_scale);
  }
  for (b in 1:nblocks) {
    sigma[b] ~ normal(sigma_loc, sigma_scale);
  }
  if (fit_model == 1) {
    for (b in 1:nblocks) {
      for (t in 1:ntrials) {
        if (y[b, t] >= 0) {
          eta[b, t] ~ normal(mulog[b], theta);
          y[b, t] ~ lognormal(eta[b, t], sigma[b]);
        }
        else {
          eta[b, t] ~ normal(log(mean(y[b, :])), 1);
        }
      }
    }
  }
}

generated quantities{
  vector<lower=0>[nblocks] expected_val;
  vector<lower=0>[nblocks] variance_val;
  matrix[nblocks, ntrials] log_lik;       // log likelihood
  for (b in 1:nblocks) {
    expected_val[b] = exp(mulog[b] + ((sigma[b] ^ 2) / 2));
    variance_val[b] = exp((2 * mulog[b]) + (sigma[b] ^ 2)) * (exp(sigma[b] ^ 2) -
1);
    for (t in 1:ntrials) {
      if (y[b, t] >= 0) {
          log_lik[b, t] = lognormal_lpdf(y[b, t] | eta[b, t], sigma[b]);
      }
    }
  }
}
```

**Figure S2, related to Figure 2, Figure 3, and Bayesian Modeling of Performance Data section:** Stan code of the Bayesian state-space model.

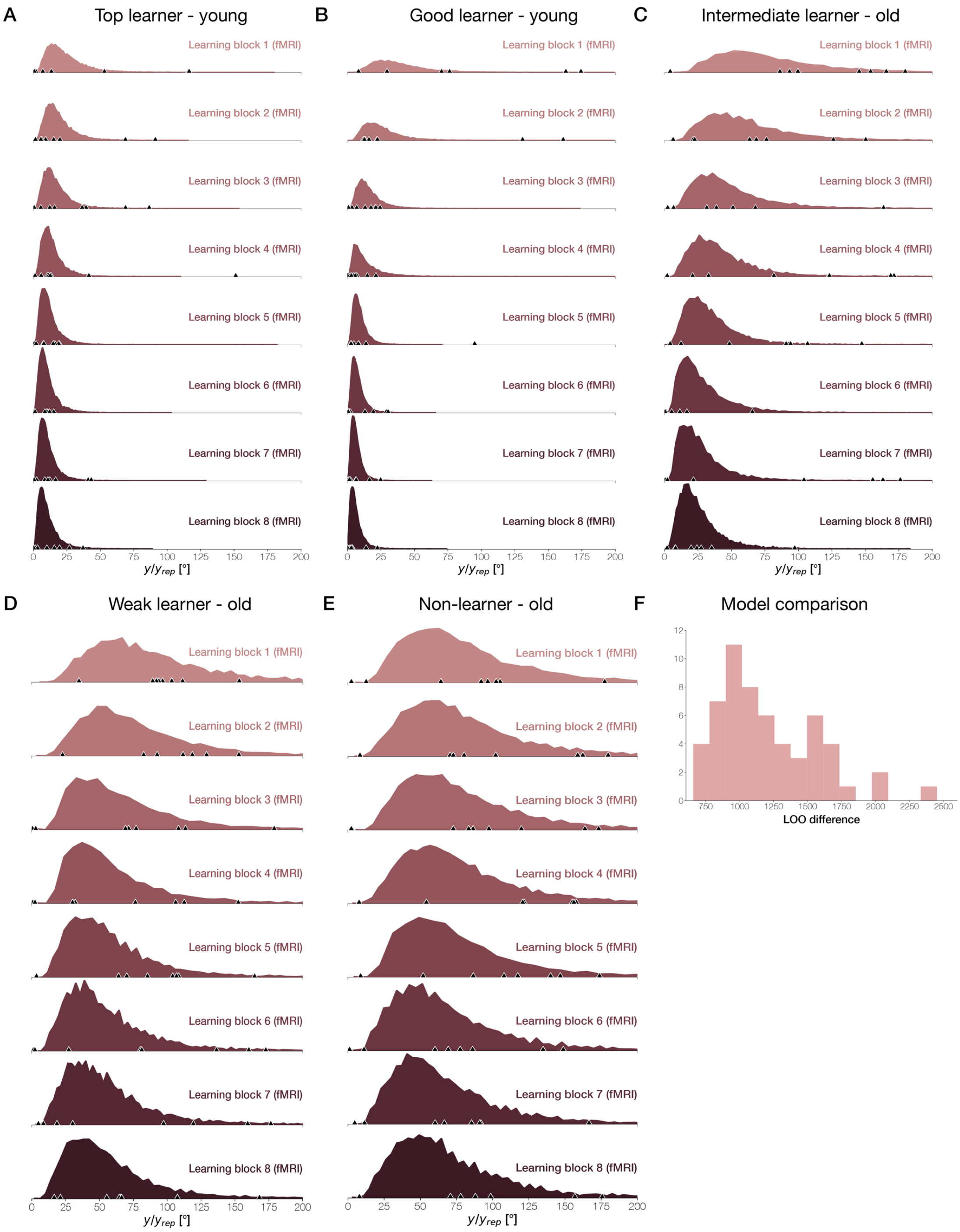


**Figure S3, related to Figure 2, Figure 3, and Bayesian Modeling of Performance Data section:** Results of the posterior predictive checks of the Bayesian state-space model for representative individuals from each learning sub-group (A: top learner young – E: non-learner old; see Performance Clustering section; the posterior predictive samples distribution, $y_{rep}$, plotted together with the observed data points, y, per learning block) and (F) histogram of the individuals' loo differences for the comparison of the Bayesian state-space model incorporating the effects of the responses, η, to an alternative model that estimated the individuals' learning state trial-wise. More positive values indicate a better fit of the first model.

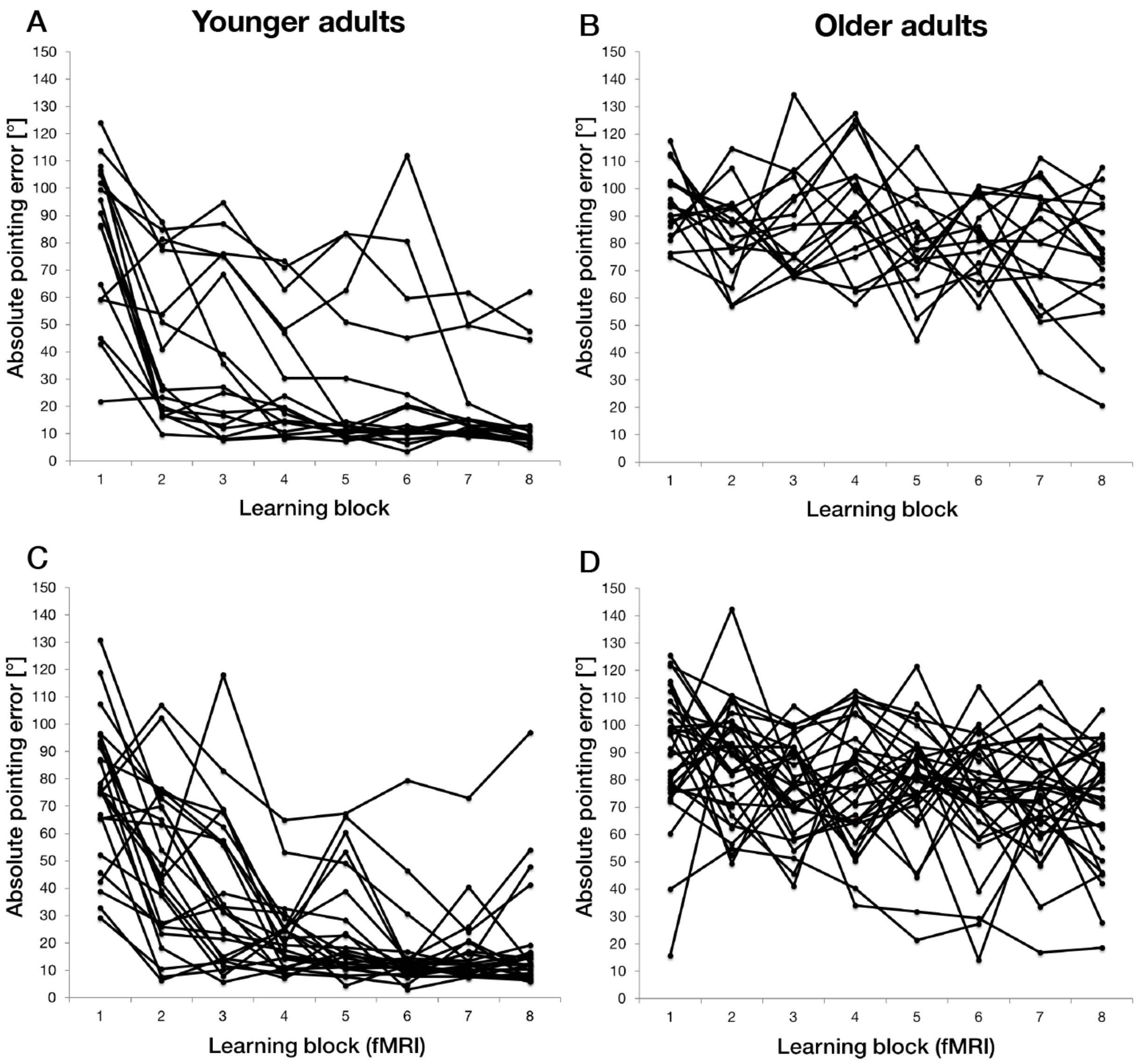


**Figure S4, related to Figure 2 and 3:** Average absolute pointing errors per learning block for each participant in (A) the younger and (B) the older age group in the behavioral experiment, and for each participant in (C) the younger and (D) the older age group in the fMRI experiment.

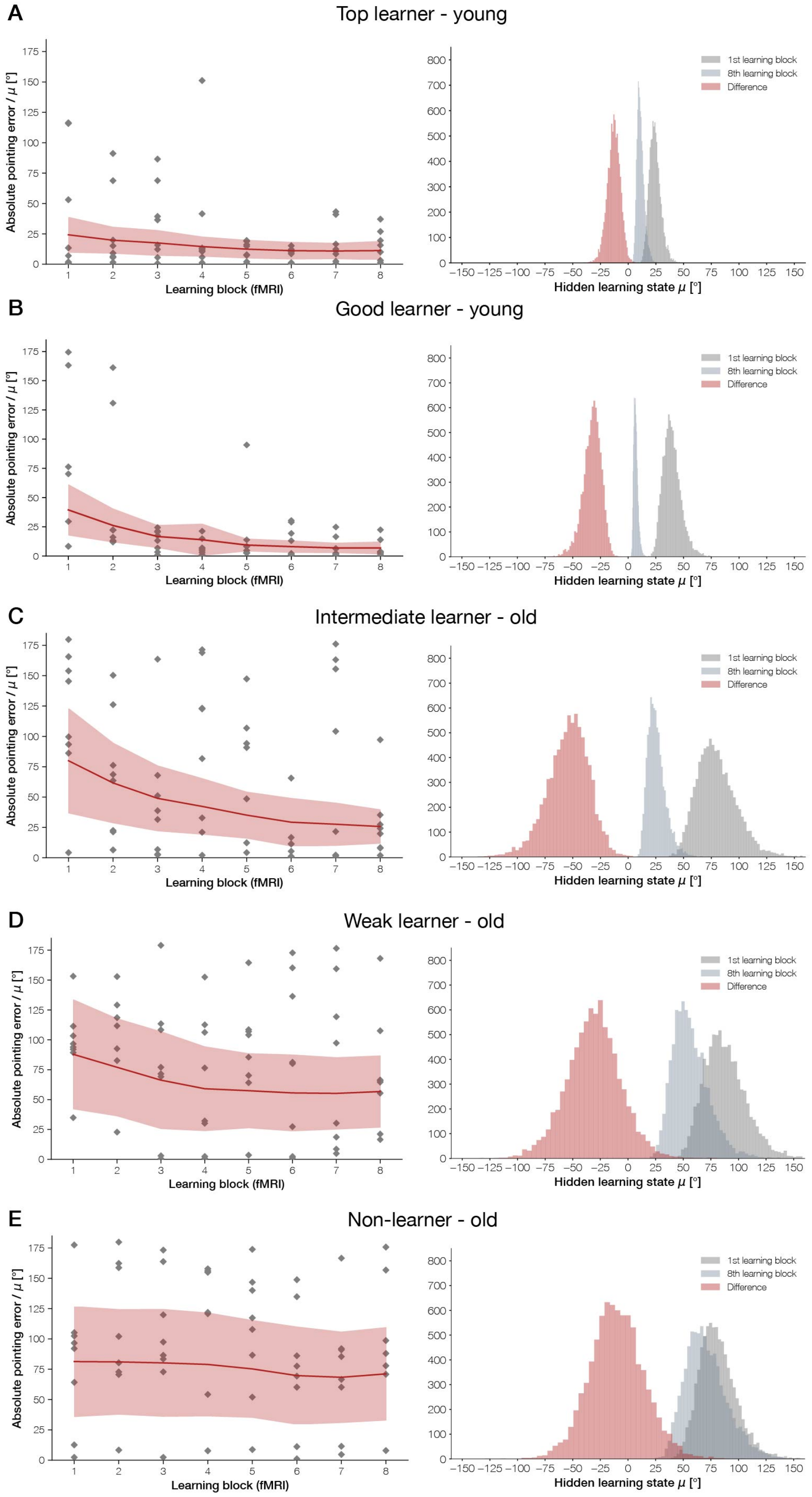


**Figure S5, related to Figure 3 and Performance Clustering section:** Definition of learning sub-groups. Difference of the latent state distributions between the last and first learning blocks (right) and their underlying hidden learning states and trial-wise performance data per learning block (left) from representative individuals from each learning sub-group in the fMRI experiment (A: top learner young – E: non-learner old).

| | Brain region | Cluster size | MNI coordinate | | | Z-score |
|---|---|---|---|---|---|---|
| | | | x | y | z | |
| A) | Navigation vs. control across the whole sample | | | | | |
| | L Parieto-occipital sulcus | 5009 | -18 | -60 | 21 | Inf |
| | R Retrosplenial cortex | | 15 | -54 | 21 | Inf |
| | L Angular gyrus | | -33 | -81 | 34 | Inf |
| | R Premotor cortex | 2594 | 33 | 6 | 57 | Inf |
| | L Premotor cortex | | -30 | 27 | 1 | 7.54 |
| | Supplementary motor area | | 0 | 15 | 47 | 7.18 |
| | R Cerebellum | 204 | 30 | -63 | -32 | 6.58 |
| | | | 39 | -69 | -29 | 6.42 |
| | L Inferior frontal gyrus | 201 | -36 | 51 | 4 | 4.91 |
| | L Middle frontal gyrus | | -36 | 54 | 18 | 4.76 |
| B) | Reduced activity in older compared to younger adults during navigation vs. control | | | | | |
| | L Cerebellum | 69 | -36 | -45 | -35 | 4.91 |
| | L Precuneus | 245 | -9 | -78 | 54 | 4.80 |
| | R Precuneus | | 12 | -69 | 57 | 4.45 |
| | L Supramarginal gyrus | 82 | -48 | -42 | 57 | 4.08 |
| | R Superior parietal lobe | 70 | 39 | -42 | 51 | 4.07 |
| | R Supramarginal gyrus | 94 | 57 | -30 | 41 | 4.01 |
| | R Postcentral gyrus | | 66 | -18 | 37 | 3.30 |
| | R Insula | 73 | 30 | 18 | 8 | 3.82 |
| | R Premotor cortex | | 48 | 6 | 14 | 3.66 |
| C) | Increased activity in older compared to younger adults during navigation vs. control | | | | | |
| | R Superior frontal gyrus | 2292 | 3 | 60 | 37 | 6.06 |
| | L Ventromedial prefrontal cortex | | -3 | 63 | 18 | 5.56 |
| | L Orbitofrontal cortex | 380 | -36 | 33 | -12 | 6.03 |
| | L Inferior frontal gyrus | | -51 | 24 | -9 | 4.54 |
| | R Middle temporal gyrus | 471 | 69 | -18 | -15 | 5.81 |
| | | | 60 | -9 | -29 | 4.84 |
| | L Cerebellum | 525 | -42 | -78 | -38 | 5.71 |
| | L Visual cortex (extrastriate area) | | -27 | -78 | -9 | 4.69 |
| | L Angular gyrus | 368 | -54 | -66 | 28 | 5.38 |
| | R Orbitofrontal gyrus | 232 | 30 | 33 | -12 | 5.32 |
| | R Inferior frontal gyrus | | 42 | 36 | -15 | 4.68 |
| | R Visual cortex (extrastriate area) | 387 | 27 | -84 | -15 | 5.13 |
| | R Cerebellum | | 15 | -87 | -32 | 4.40 |
| | R Angular gyrus | 245 | 45 | -57 | 31 | 4.99 |
| | | | 54 | -66 | 47 | 4.82 |
| | L Superior temporal sulcus | 487 | -63 | -39 | 1 | 4.84 |
| | | | -57 | -3 | -15 | 4.35 |
| | L Hippocampus | 82 | -21 | -18 | -12 | 4.76 |
| | | | -30 | -15 | -22 | 4.21 |

| | | | | | | |
|---|---|---|---|---|---|---|
| | Posterior cingulate cortex | 192 | 0 | -60 | 34 | 4.57 |
| | L Posterior cingulate cortex | | -6 | -57 | 21 | 4.32 |
| | L Premotor cortex | 354 | -39 | 12 | 24 | 4.55 |
| | L Middle frontal gyrus | | -42 | 21 | 47 | 4.26 |
| | R Precentral gyrus | 120 | 33 | -15 | 28 | 4.23 |
| | | | 36 | -15 | 41 | 4.00 |
| D) | Retrieval vs. encoding across the whole sample | | | | | |
| | R Caudate | 8550 | 12 | 12 | -2 | Inf |
| | L Premotor Cortex | | -48 | 0 | 34 | Inf |
| | L Fusiform Gyrus/Inferior temporal gyrus | | -42 | -63 | -12 | Inf |
| | L Parieto-occipital sulcus | | -15 | -72 | 24 | 7.81 |
| | L Lingual Gyrus | | -21 | -54 | -2 | 7.79 |
| | R Cerebellum | 241 | 6 | -69 | -22 | 5.73 |
| E) | Encoding vs. retrieval across the whole sample | | | | | |
| | L Visual cortex (extrastriate area) | 9128 | -6 | -87 | -9 | Inf |
| | R Visual cortex (extrastriate area) | | 12 | -90 | -9 | Inf |
| | R Superior frontal gyrus | 387 | 30 | 39 | 51 | 4.89 |
| | | | 39 | 30 | 41 | 4.78 |
| | L Superior frontal gyrus | 118 | -36 | 30 | 51 | 4.78 |
| | | | -24 | 27 | 51 | 4.00 |
| | L Superior frontal gyrus | 101 | -18 | 57 | 31 | 3.42 |
| | | | -9 | 54 | 31 | 3.41 |
| | R Anterior cingulate cortex | 86 | 3 | 21 | 24 | 4.15 |
| F) | Reduced activity in older compared to younger adults during retrieval vs. encoding | | | | | |
| | R Fusiform gyrus | 144 | 27 | -63 | -12 | 4.56 |
| | | | 36 | -78 | -15 | 4.17 |
| | L Brain stem | 92 | -3 | -45 | -48 | 4.17 |
| | | | 0 | -39 | -38 | 3.81 |
| G) | Increased activity in older compared to younger adults during retrieval vs. encoding | | | | | |
| | R Supramarginal gyrus | 2863 | 63 | -24 | 24 | 6.23 |
| | R Inferior frontal gyrus | | 57 | 15 | -2 | 5.16 |
| | L Postcentral gyrus | 196 | -24 | -39 | 67 | 5.61 |
| | R Posterior cingulate cortex | 438 | 6 | -27 | 41 | 5.56 |
| | | | 9 | -51 | 34 | 4.07 |
| | L Middle temporal gyrus | 2422 | -54 | -54 | -2 | 5.44 |
| | L Insula | | -39 | -15 | -5 | 5.43 |
| | L Superior temporal gyrus | | -66 | -39 | 24 | 5.40 |
| | R Rostral prefrontal cortex | 1180 | 9 | 54 | -2 | 4.59 |
| | L Anterior cingulate cortex | | -3 | 27 | 18 | 4.50 |
| | R Superior frontal gyrus | | 24 | 51 | 34 | 4.42 |
| | L Cerebellum | 220 | -27 | -84 | -32 | 4.49 |
| | | | -39 | -72 | -42 | 4.22 |
| | L Superior frontal gyrus | 99 | -27 | 45 | 24 | 4.07 |

| | | | | | |
|---|---|---|---|---|---|
| L Middle frontal gyrus | | -33 | 57 | 21 | 3.62 |
| H) Age-group differences in learning-related activity decreases | | | | | |
| L Ventromedial prefrontal cortex | 202 | -12 | 54 | -2 | 4.04 |
| | | -3 | 42 | -12 | 4.02 |
| R Superior frontal gyrus | 71 | 0 | 42 | 37 | 4.02 |
| | | 9 | 45 | 41 | 3.82 |
| I) Age-group differences in learning-related activity increases | | | | | |
| L Parieto-occipital sulcus | 2257 | -9 | -66 | 24 | 5.93 |
| | | -27 | -69 | 51 | 5.52 |
| R Retrosplenial cortex | | 9 | -57 | 4 | 4.82 |
| Supplementary motor area | 203 | 3 | 3 | 61 | 5.45 |
| | | -9 | 6 | 41 | 3.81 |
| R Premotor cortex | 125 | 30 | 0 | 51 | 4.81 |
| | | 27 | -3 | 67 | 4.44 |
| L Premotor cortex | 138 | -24 | 3 | 57 | 4.57 |
| | | -30 | -6 | 57 | 4.49 |
| Visual cortex (striate area) | 257 | -6 | -102 | -2 | 4.54 |
| | | 12 | -102 | 11 | 3.96 |
| J) Learning-group differences in learning-related activity changes within older adults | | | | | |
| R Lateral frontopolar cortex | 117 | 33 | 48 | -9 | 4.84 |
| R Ventrolateral prefrontal cortex | | 39 | 36 | -5 | 4.75 |
| | | 51 | 36 | -5 | 4.62 |
| R Insula | 76 | 33 | 18 | -12 | 4.54 |
| | | 33 | 18 | 1 | 4.15 |

**Table S1, related to Table 1, Figure 4, and Figure 5:** Spatial coordinates of the local maxima in the whole-brain fMRI analyses ($p < 0.05$, whole-brain FWE-corr.).

**Data S1, related to Figure 3**

*Additional analyses in the behavioral experiment*

Age-related differences in pointing performance depending on the characteristics of the VE were further analyzed by means of an ANOVA on the absolute pointing errors with intersection (I1-I4), direction (D1-D4), and target landmark (town hall, church) as repeated measures variables and age group (younger adults, older adults) as between-subjects variable. A significant interaction between the four factors suggested that the performance of the age groups was modulated by the respective intersection-direction-target landmark combination encountered in the VE during retrieval, $F(9, 288) = 2.05$, $p = .034$. Therefore, follow-up ANOVAs were conducted within the two age groups separately. In younger adults, a significant main effect of intersection, $F(3, 48) = 6.18$, $p = .005$ (Greenhouse-Geißer corrected), showed that performance was worse when they were located at I4 ($M = 48.6° \pm 21.9°$) as compared to I1 ($M = 25.0° \pm 19.7°$) or I2 ($M = 30.6° \pm 25.7°$), all $p \leq .010$ (Bonferroni-corrected). This was modulated by a significant interaction between intersection and target landmark, $F(3, 48) = 11.4$, $p < .001$. Pointing errors were smaller in this age group when they pointed towards the town hall ($M = 13.5° \pm 14.5°$) as compared to the church ($M = 36.6° \pm 29.3°$) at I1, which was the intersection adjacent to the town hall, and vice versa at I3, which was the intersection adjacent to the church (town hall: $M = 45.5° \pm 35.5°$; church: $M = 25.8° \pm 26.9°$), all $t \geq 3.33$, all $p \leq .004$. The directions from which the intersections were approached did not seem to have an influence on performance in this age group. In older adults, the same ANOVA also revealed an interaction between intersection and target landmark, $F(3, 48) = 3.38$, $p = .026$. When located at I3, pointing errors were smaller when the target landmark was the adjacent church ($M = 73.2° \pm 27.4°$) as compared to the town hall ($M = 94.2° \pm 19.9°$), $t(16) = 2.73$, $p = .015$. The corresponding comparison for I1 did not reach significance, $t(16) = 1.12$, $p = .281$. In addition, there was a significant interaction between direction and target landmark, $F(3, 48) = 3.75$, $p = .039$ (Greenhouse-Geißer corrected). Post-hoc t-tests showed that pointing towards the town hall ($M = 80.3° \pm 18.1°$) tended to be easier as compared to pointing towards the church ($M = 94.5° \pm 27.1°$) for the older adults when they approached the intersections from D4 (i.e., facing east), $t(16) = 1.96$, $p = .068$. In contrast, pointing towards the church ($M = 69.9° \pm 19.6°$) tended to be easier than pointing towards the town hall ($M = 87.0° \pm 23.3°$) when they approached the intersections from D2 (i.e., facing west), $t(16) = 1.99$, $p = .064$. This was modulated by an interaction between intersection, direction, and target landmark, $F(9, 144) = 2.25$, $p = .022$. Separate follow-up ANOVAs for each intersection with direction (D1-D4) and target landmark (town hall, church) as repeated measures variables revealed for I1 an interaction between the two factors that approached significance, $F(3, 48) = 3.21$, $p = .067$ (Greenhouse-Geißer corrected). Pointing towards the adjacent town hall ($M = 56.5° \pm 41.3°$) was easier than pointing towards the church ($M = 96.9° \pm 38.1°$) when participants were coming from D4 (i.e., facing east), $t(16) = 3.54$, $p = .003$. At I2, a main effect of direction, $F(3, 48) = 5.25$, $p = .003$, indicated that pointing generally tended to be easier from D2 (i.e., facing west; $M = 68.8° \pm 32.7°$) as compared to D1 ($M = 94.0° \pm 24.5°$) or D4 ($M = 95.8° \pm 37.1°$), that is, when they were facing towards the deadends at this intersection, all $p \leq .054$ (Bonferroni-corrected). At I3, a main effect of target landmark indicated that pointing towards the adjacent church (M = 73.2° ± 27.4°) was easier for the older adults than pointing towards the town hall (M = 94.2° ± 20.0°), $F(3, 48) = 7.47$, $p = .015$. This was modulated by an interaction between direction and target landmark, $F(3, 48) = 4.44$, $p = .008$. Pointing towards the church was easier when coming from D1 (i.e., facing south; town hall: M = 107.7° ± 31.5°; church: M = 65.6° ± 36.0°) or D2 (i.e., facing west; town hall: M = 98.0° ± 39.4°; church: M = 52.6° ± 40.8°), all $t \geq 3.19$, all $p \leq .006$. Finally, at I4, there was also an interaction between direction and target landmark, $F(3, 48) = 3.74$, $p = .017$. Performance was better when participants pointed towards the church (M = 75.9° ± 30.8°) as compared to the town hall (M = 106.6° ± 31.4°) when approaching the intersection from D2 (i.e., facing west), $t(16) = 2.59$, $p = .020$.

Taken together, this analysis supports the results from the analyses reported in the main text implying better navigational encoding in younger adults and a higher reliance on the specific sensory input in older adults. The directions from which they were approaching the intersections partly seemed to have an impact on their performance although variability in performance was generally quite high in this age group. One should note that a corresponding analysis restricted to the last learning blocks where participants were better able to navigate was not conducted due to a lack of statistical power. This would have also resulted in unequal numbers of trials that enter the analysis given the number of trials per condition and their randomization across the entire experiment.
